# Immunosuppressive tumor microenvironment of osteosarcoma

**DOI:** 10.1101/2023.11.01.565008

**Authors:** Aaron Michael Taylor, Jianting Sheng, Patrick Kwok Shing Ng, Jeffrey M. Harder, Parveen Kumar, Ju Young Ahn, Yuliang Cao, Alissa M. Dzis, Nathaniel L. Jillette, Andrew Goodspeed, Avery Bodlak, Qian Wu, Michael S. Isakoff, Joshy George, Jessica D.S. Grassmann, Diane Luo, William F. Flynn, Elise T. Courtois, Paul Robson, Masanori Hayashi, Alini Trujillo Paolillo, Antonio Sergio Petrilli, Silvia Regina Caminada de Toledo, Fabiola Sara Balarezo, Adam D. Lindsay, Bang Hoang, Stephen T.C. Wong, Ching C. Lau

**Affiliations:** The Jackson Laboratory for Genomic Medicine, Farmington, CT 06032; Systems Medicine and Bioengineering Department, Houston Methodist Neal Cancer Center, Houston, TX 77030; The Jackson Laboratory, Bar Harbor, ME 04609; Department of Biomedical Engineering, Texas A&M University, College Station, TX, USA; University of Colorado Anschutz Medical Campus, Aurora, CO 80045; University of Connecticut School of Medicine, Farmington, CT 06032; Connecticut Children’s Medical Center, Hartford, CT 06106; Children’s Hospital Colorado, Aurora, CO 80045; Hospital do GRAACC, São Paulo, Brazil; Hartford Hospital, Hartford, CT 06106; Montefiore Medical Center, Bronx, NY 10467; Departments of Radiology, Neuroscience, Pathology and Laboratory Medicine, Weill Cornell Medicine, Cornell University NY, NY, USA

## Abstract

Osteosarcoma is the most common malignant bone tumor in children, characterized by a high degree of genomic instability, resulting in copy-number alterations and genomic rearrangements without disease-defining recurrent mutations. Clinical trials based on molecular characterization have failed to find new effective therapies or improve outcomes over the last 40 years. To better understand the immune microenvironment of osteosarcoma, we performed single-cell RNA sequencing on six tumor biopsy samples, combined with a previously-published cohort of six samples. Additional osteosarcoma samples were profiled using spatial transcriptomics for validation of discovered subtypes and to add spatial context. Analysis revealed immunosuppressive cells, including myeloid-derived suppressor cells (MDSCs), regulatory and exhausted T-cells, and LAMP3+ dendritic cells. Using cell-cell communication modeling, we identified robust interactions between MDSCs and other cells, leading to NF-κB upregulation and an immunosuppressive microenvironment, as well as interactions involving regulatory T-cells and osteosarcoma cells that promoted tumor progression and a proangiogenic niche.

**Statement of Significance:** Osteosarcoma patient survival has remained stagnant for several decades due to lack of successful therapy for high-risk patients, including those with metastatic disease. Identifying novel therapeutics including immunotherapies is of great clinical importance. Our study highlights several important immunosuppressive mechanisms within osteosarcoma that should be considered when developing future immunotherapies.

## Introduction

Osteosarcoma (OS) is the most common primary malignant bone tumor affecting children and young adults. Current treatment with chemotherapy and surgical resection results in a long-term overall survival rate of ∼70% for patients with localized disease but only 20-30% for patients with metastatic disease (1). No significant advances in treatment or overall survival rate have been made in the last four decades.

Genomic profiling studies of OS tumors from predominantly pediatric populations have shown high levels of chromosomal structural variation, including massive rearrangements resulting from chromothripsis and hypermutation in localized regions (kataegis) that contribute to significant tumoral heterogeneity (2). Because of their potential to generate novel rearrangements in protein-coding sequences, the genomic structural abnormalities observed in OS would be expected to express neoantigens that should serve as potent targets for immunotherapy (IT). Aligning with that, a study of 48 OS patients demonstrated that PD-L1 expression was found in 25% of primary OS samples and correlated with immune cell infiltration and event-free survival (3). Anti-PD-L1 and anti-CTLA4 treatment also controlled OS tumor growth in a mouse model (4). However, the clinical trials using anti-PD-L1 and anti-CTLA4 have not shown any efficacy in OS treatment (5,6).

The tumor microenvironment (TME) of multiple cancers has been shown to play a critical role in tumor progression and treatment response (7). Previous studies of the OS TME have revealed the presence of multiple immune cell types, e.g., dendritic cells (DCs), macrophages, neutrophils, and lymphoid cells, with a wide range of cell abundance across patient samples (8–10). However, the role of these tumor-infiltrating immune cells in OS is not fully understood. With the recent advancements in single-cell technologies, several groups have applied single-cell RNA sequencing (scRNA-seq) analysis on limited populations of OS patient tumors, including biopsy, post-treatment surgical resection, and lung metastasis samples (11,12). Apart from the heterogeneous OS tumor cells, those studies also identified several molecular subtypes of immune cells, including *TIGIT*+ exhausted T cells, *FOX3P*+ regulatory T cells, and *LAMP3*+ mature regulatory DCs (11–13). Taken together with the failure of the IT clinical trials, the complex TME within OS may serve an immunosuppressive function. Thus, a better understanding of the OS TME is urgently needed to improve the IT efficacy for OS patients.

To advance our understanding of the OS TME naïve to chemotherapy-induced biological changes, we performed scRNA-seq analysis on six pre-treatment primary tumor biopsy samples of OS. We combined our data with the published six pre-treatment OS sample data (11) to increase our ability to detect rare cell types or subtypes. We described a number of immune cells that potentially contribute to the tumor progression and immunosuppressive TME using subclustering, differential gene expression, and pathway analysis. In addition, we evaluated the spatial distribution of those cell types of interest by spatial transcriptomics analysis on additional pre-treatment biopsy samples of OS.

## Results

### Single-cell transcriptomic analysis of treatment-naïve osteosarcoma

The tumor microenvironment (TME) plays a critical role in tumor progression and treatment response (14–16). Chemotherapy, as the standard treatment regimen for OS, will change the composition of the TME. To identify the TME cellular composition and understand the role of different cell types on tumor progression and treatment response without the influence of treatment, we aimed to investigate the TME of treatment-naïve OS tumors. We carried out scRNA-seq of six treatment-naïve OS tumors (**Supplementary Table S1**) using the 10X Genomics Chromium platform. After quality control and doublet removal, we obtained a dataset of 22,035 cells from six OS tumors (**Supplementary Table S1**). The number of detected UMIs ranged from 694 to 155,389 per cell, with the number of detected genes ranging from 500 to 8,709. To increase our power to identify rare cell types or subtypes, we leveraged the published scRNA-seq dataset from another six treatment-naïve tumors (GSE162454). A total of 40,588 additional cells passed our internal quality control filters and were subsequently integrated with our dataset using the *Harmony* package. A combined dataset with a total 62,623 cells from 12 treatment-naïve OS tumors was generated. New data generated from our scRNA-seq cohort contributes 35.19% of cells to the combined dataset (**Supplementary Table S1, Supplementary Table S2**).

To determine the cellular composition of the tumors, we performed unsupervised clustering via *Seurat*’s graph-based clustering method. Seventeen major clusters were identified (**Figure 1A**). Cell type identification of each cluster was made based on the cluster-specific differential gene expression (**Supplementary Table S3**) and literature-based cell type markers. The expression of each marker gene was shown in **Figure 1D**. We identified two clusters of macrophages/DCs (*MSR1*, *C1QC*, *FOLR2*), six clusters of osteoblastic tumor cells (*SATB2*, *IBSL*, *ALPL*), one cluster of natural killer (NK) and T-cells (*CD3D, NKG7 and TRBC1*), osteoclasts (*ACP5*, *CTSK*, *MMP9*), monocytes/neutrophils (*S100A8*, *S100A9*, *FCN1*), fibroblasts (*TAGLN*, *ACTA2*, *FAP*), B-cells (*MS4A1*, *CD79A*, *BANK1*), plasma cells (*IGHG1*, *IGLC2*, *IGHG4*), mast cells (*TPSB2*, *TPSAB1*, *CPA3*) and endothelial cells (*CLEC14A*, *PLVAP*, *VWF*). The smallest cluster contained only 36 cells (0.06% of total cells) with no known cell type marker identified, and poor total gene and UMI counts compared to the other clusters. It was assigned as a cluster comprised of low-quality cells and removed prior to downstream analyses. Except for the low-quality cluster and one osteoblastic tumor cell cluster, each cluster consisted of cells from both published and our new datasets (**Figure 1B, Supplementary Figure 1**). The proportion of each cell type and distribution of cells in UMAP is similar between the published and new datasets (**Figure 1B** and **1C**). Since the cells in each significant cluster also expressed well-known, cell-type-specific genes, merging the two datasets was successful.

**Figure 1:**
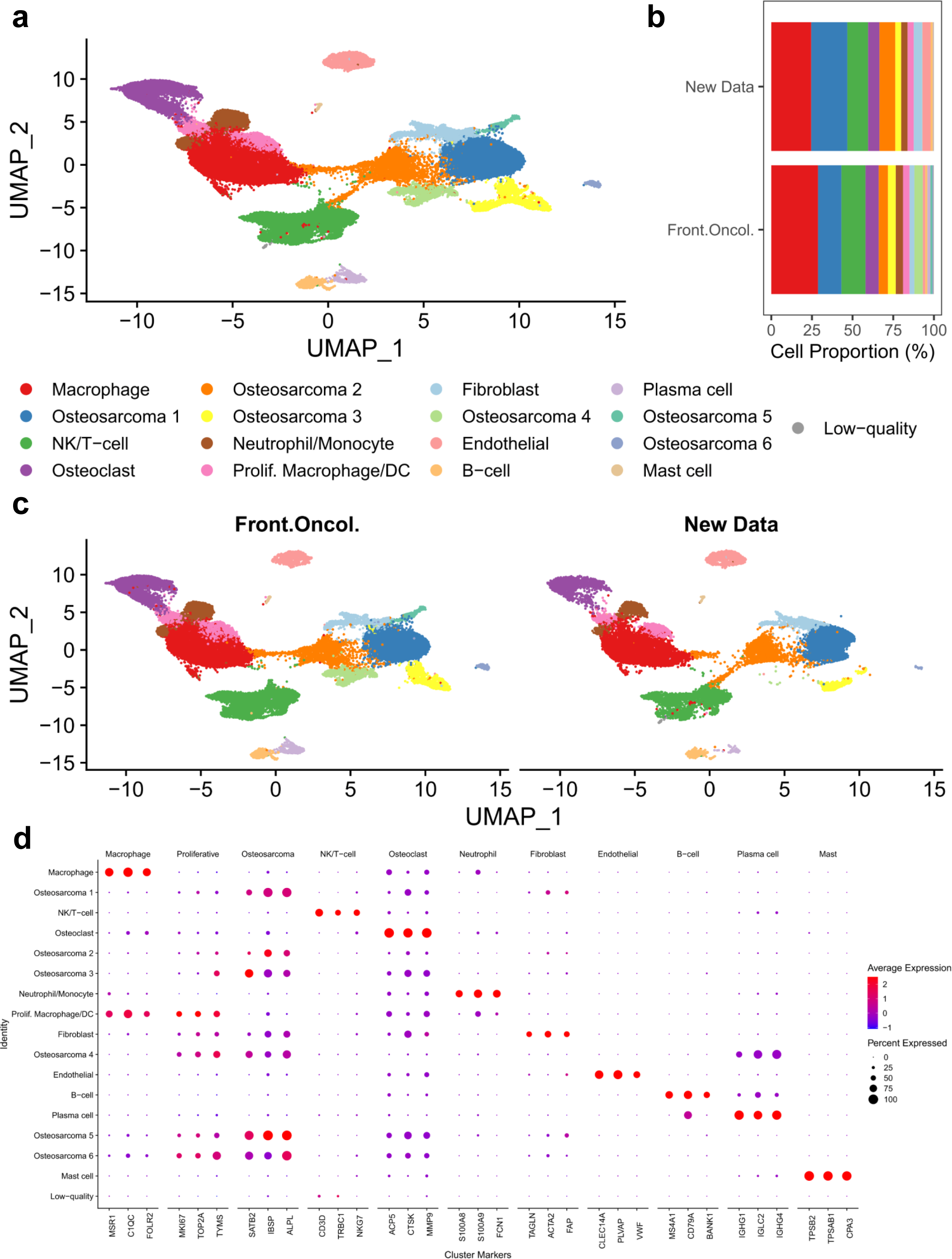
Primary clustering identifies major cell types in pre-treatment osteosarcoma tumors. **(A)** UMAP visualization of *Harmony*-corrected principal components, with cell type clusters separated by color. **(B)** Cell type proportions of each sample cohort. Macrophages and osteosarcoma were the most abundant cell types in both cohorts. **(C)** UMAP visualization of *Harmony*-corrected principal components in each cohort. **(D)** Expression of selected cell-type-specific markers in major cell clusters. Markers were chosen from the differentially expressed genes in **Supplemental Table 2** based on fold-change, cell cluster specificity, and their prior published use as cell type markers.

### Macrophages and dendritic cells

The roles of different immune cells in the TME can be diverse and highly dependent on their phenotypic status. To understand the role of different subtypes of immune cells in OS, we looked further into the phenotypic subtypes of various immune cells. First, we investigate the most abundant cell lineage – myeloid cells. Two major myeloid cell clusters were identified in the present dataset – a predominantly macrophage cluster and a macrophage/DC cluster that expressed proliferative markers. While both clusters have strong expression of macrophage markers (*MSR1*, *C1QC*, *FOLR2*), the proliferative macrophage/DC cluster also has expression of cell proliferation genes (*MKI67*, *TOP2A*, *TYMS*) (**Figure 1D**). Further unsupervised clustering of the macrophage cluster revealed seven sub-clusters with distinct differentially expressed genes (**Supplementary Table S4**). We were able to detect classically activated macrophages (*NR4A3*, *IL1B*, *CCL3/4*, *APOE*, *TXNIP*), *LYVE*+ macrophages, interferon-stimulated macrophages (*CXCL10*, *ISG15*, *IFIT1*), angiogenic macrophages (*SPP1*, *ADAM8*, *VIM*, *VCAN*) and one macrophage cluster expressing T-cell maker genes (*IL32*, *NKG7*, *TRAC*) (**Figure 2A** and **2B**).

**Figure 2:**
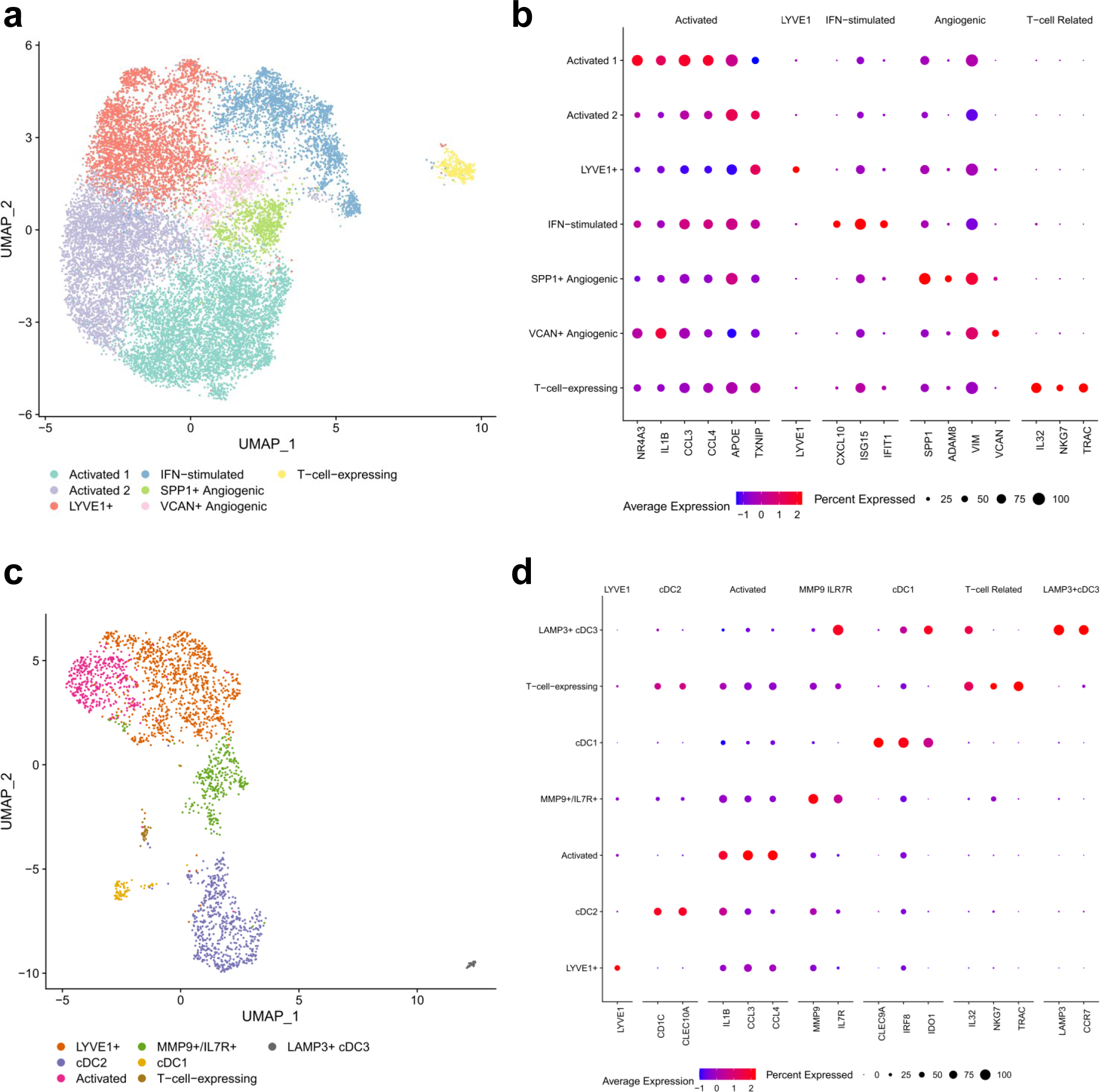
Subclustering of macrophage/DC clusters **(A)** UMAP visualization of *Harmony*-corrected principal components of the macrophage cluster. Cell subtypes separated by color. **(B)** Expression of selected cell-subtype-specific markers amongst macrophage subclusters. Several subtypes of activated, interferon (IFN)-stimulated, and angiogenic macrophages were identified. **(C)** UMAP visualization of *Harmony*-corrected principal components of the proliferative macrophage/DC cluster. Cell subtypes separated by color. **(D)** Expression of selected cell-subtype-specific markers amongst macrophage/DC subclusters. *LYVE*+ macrophages were detected, along with populations of cDC1 and cDC2, in addition to a putative *LAMP3*+ mregDC population.

Although the role of various phenotypic macrophages in OS is not fully elucidated yet, their contribution to the anti-inflammatory and immunosuppressive TME in OS has been suggested based on literature. The *LYVE*+ macrophage subpopulation has been found in multiple human tissues as tissue-resident macrophages and has been demonstrated to support angiogenesis in different tissues (17–21). In cancer, they have been shown to cooperate with mesenchymal cells in the TME and support tumor growth in a mammary adenocarcinoma model (22). Two subtypes of macrophages with angiogenesis signatures (*SPP1+*, *ADAM8+*, and *VIM+*, or *VCAN+, respectively*) were also detected. *SPP1*+ and *VCAN*+ angiogenic macrophages have been associated with worse clinical outcomes in multiple types of epithelial cancers (23). The *SPP1*+ macrophage cluster also expressed *HMOX1*, which has been shown to play a role in the immunosuppressive program of tumor-associated macrophages (TAM) (24). The *VCAN*+ angiogenic macrophages have high expression of epidermal growth factor (EGF) family genes, including *EREG* and *AREG*. M2-like TAM-secreted EGF has been shown to promote metastasis in ovarian cancer (25). Interestingly, this population of macrophages also has higher expression of *OLR1*, a marker of myeloid-derived suppressor cells (MDSC) in other tumors (26). Macrophages expressing T-cell marker genes have been previously found in inflammatory diseases and head and neck squamous cell carcinoma (27,28), however, their role in the OS TME is unclear.

Subclustering of the proliferative macrophage/DC cluster revealed seven subpopulations, including three subclusters of dendritic cells and four macrophage subgroups (**Supplementary Table S5**). Classic dendritic cells 1 (cDC1s) and cDC2s were identified based on the expression of *CLEC9A*, *IRF8* and *IDO1* and *CD1C* and *CLEC10A*, respectively (**Figure 2C** and **2D**). We also detected another distinct type of dendritic cells based on high expression of *LAMP3*, *CCR7*, and *IDO1*. The cDC2 population was found to be more abundant than cDC1 and *LAMP3*+ DCs in the tumors studied. The presence of *LAMP3*+ dendritic cells (*LAMP3*+ cDCs) or mature regulatory dendritic cells (mregDCs) has also been reported recently in various cancer types, including osteosarcoma (13,23). One possible mechanism of mregDCs in immunosuppression in OS TME has been suggested through interaction with regulatory T-cells (T-regs) via *CD274-PDCD1* and *PVR-TIGIT* signaling (13).

Among proliferative macrophages, similar macrophage subtypes were found as discussed previously, including *CCL3-4*+/*IL1B*+ classically-activated macrophages, *SSP1*+/*LYVE*1+ angiogenic macrophages, and macrophages expressing T-cell associated genes (*IL32+*, *NKG7+, TRAC+*). A population of *MMP9*+/*IL7R*+ macrophages was found only within the proliferative macrophage cluster.

### Immunosuppressive neutrophils/Myeloid-derived suppressor cells (MDSC)

Neutrophils have been known to promote tumor growth, angiogenesis, metastasis and inhibit anti-cancer T-cell activity (29). To investigate the potential immunosuppressive features of neutrophils in OS, subclustering of the monocyte/neutrophil cluster was carried out, which revealed four different cell types, including two populations of *S100A8/9/12*+ neutrophils, and one population of each of *CDKN1C*+/*FCGR3A*+ non-classical monocytes and *FN1*+/*SPP1*+ monocyte-like cells (**Figure 3A** and **3B, Supplemental Table S6**). Of note, among the two neutrophil populations, the larger population of neutrophils expressed higher levels of *S100A8/9/12*, and genes previously identified as markers of myeloid-derived suppressor cells (MDSC), including *VCAN*, *CLEC4E*, and *CSF3R* (**Figure 3C**) (30). These findings suggest the presence of immunosuppressive neutrophils or MDSC in OS TME and could be one of the major contributors to immunosuppression in OS. In addition, monocyte-like cells are a possible precursor of TAM and can contribute to the accumulation of MDSCs in cancer (31).

**Figure 3:**
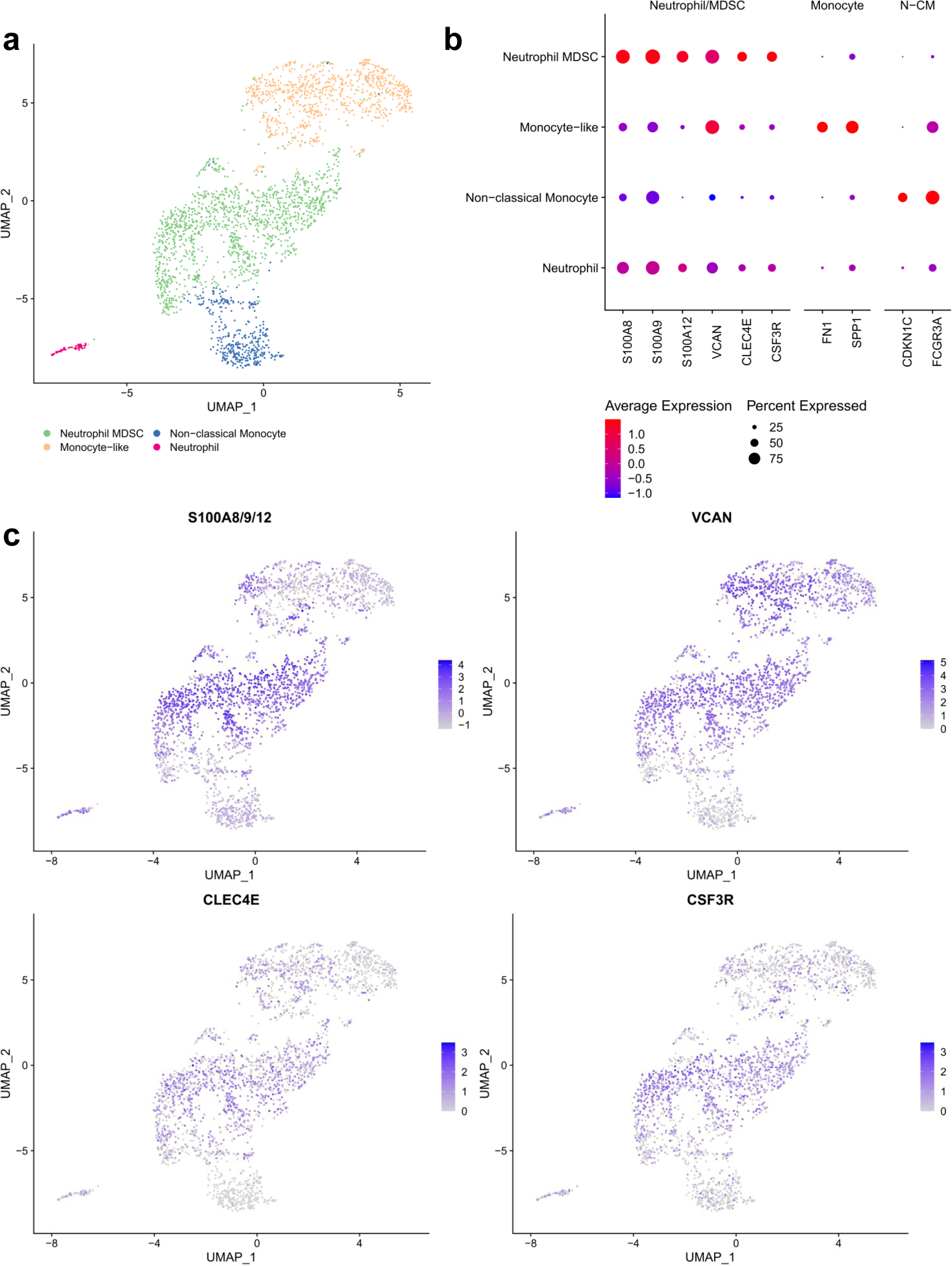
Identification of MDSCs amongst neutrophil/monocyte population **(A)** UMAP visualization of *Harmony*-corrected principal components of major neutrophil/monocyte cluster. Cell subtypes separated by color. **(B)** Dotplot expression of selected markers amongst neutrophil and monocyte subclusters (“N-CM” = non-classical monocyte). **(C)** Visualization of MDSC feature expression. *Seurat*’s “AddModuleScore” function was used to create an S100A gene expression score in the leftmost plot. Expression of other MDSC signature genes (*VCAN*, *CLEC4E* and *CSF3R*) is also shown.

### Regulatory and Exhausted T-cells

T-cells are essential for the immune anti-tumor response and for the successful administration of immunotherapy (32). Negative regulation of T-cells, leading to hypofunctional or exhausted T-cells, is a hallmark of many cancers, and understanding the mechanisms that lead to this exhaustion can provide potential targets for immunotherapy development (33,34). In order to better understand the T-cell and NK cell subtypes present within the TME of OS, we mapped cells of the NK/T cell cluster (C02) to the peripheral blood mononuclear cell (PBMC) reference dataset (35) using *Azimuth* (36). We separated populations of NK cells (*NKG7*, *KLRD1*, *TYROBP*, *GNLY*, *FCER1G*, *PRF1*, *CD247*, *KLRF1*, *CST7* and *GZMB*), helper *CD4*+ T-cells (*IL7R*, *MAL*, *LTB*, *CD4*, *LDHB*, *TPT1*, *TRAC*, *TMSB10*, *CD3D* and *CD3G*) and cytotoxic *CD8*+ T-cells (*CD8B*, *CD8A*, *CD3D*, *TMSB10*, *HCST*, *CD3G*, *LINC02446*, *CTSW*, *CD3E* and *TRAC*) (**Figure 4A** and **4B, Supplementary Table S7**).

**Figure 4:**
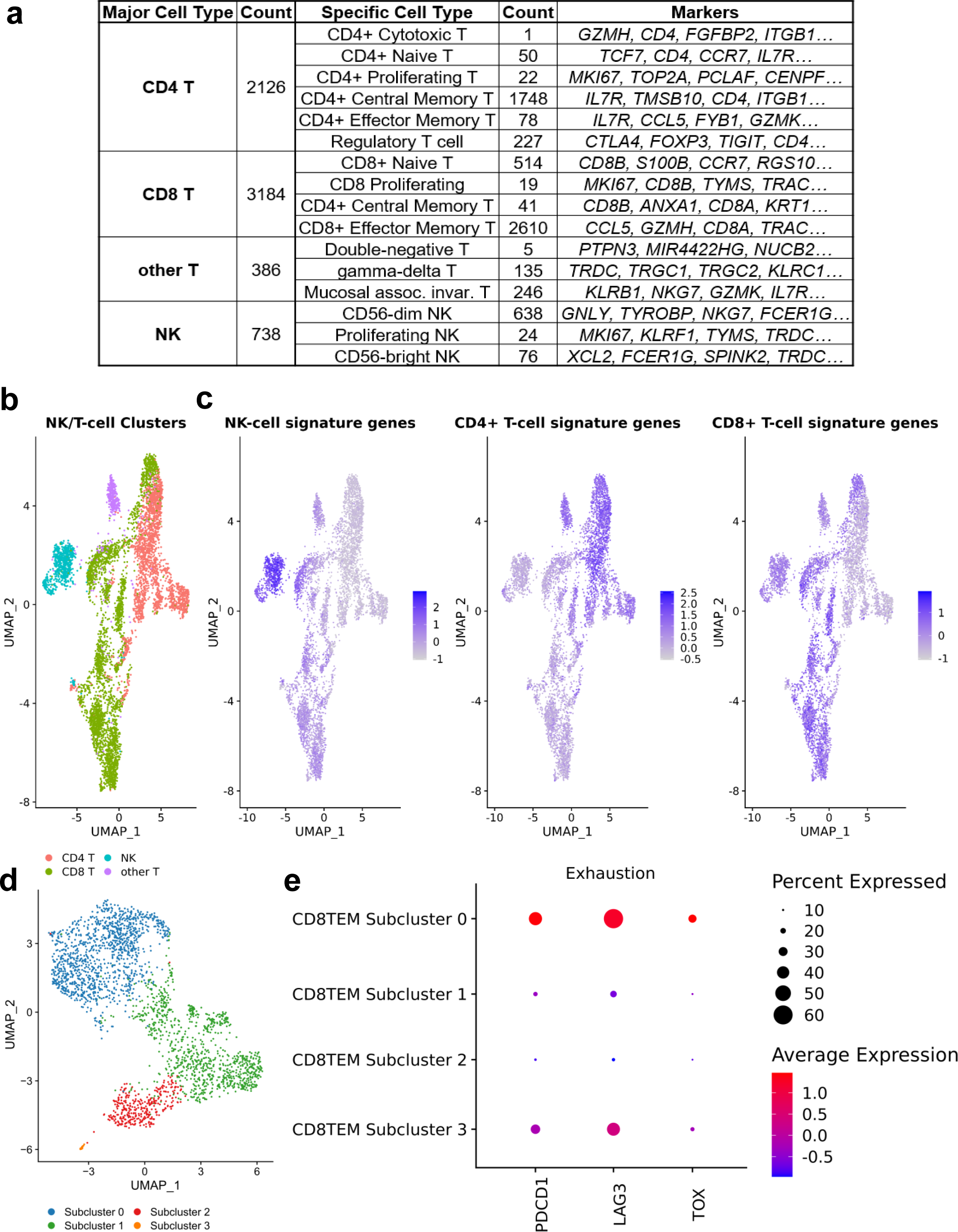
Identification of NK/T-cell subtypes using *Azimuth* **(A)** Identified NK/T-cells and their subtypes. Cells identified with >0.75 confidence score were mapped to *Azimuth*’s “L1” identities (‘Major Cell Cluster’) from the reference “PBMC” dataset. Further annotation using the “L2” cell identities (‘Specific Cell Type’) is also shown, along with a subset of the signature genes that define those cell subtypes. **(B)** UMAP visualization of *Harmony*-corrected principal components of identified NK/T-cell populations, colored by L1 identities. **(C)** NK cell, CD4+ T-cell, and CD8+ T-cell scores were created using *Seurat*’s “AddModuleScore” function from the *Azimuth*-defined cell type markers. **(D)** UMAP visualization of *Harmony*-corrected principal components of identified CD8 T-effector-memory (CD8TEM) cell subpopulations. **(E)** Dotplot expression of exhaustion markers amongst CD8TEM subpopulations. Exhaustion markers were overexpressed in the largest subcluster of CD8TEMs.

We subdivided these categories further using *Azimuth* to detect the major subtypes of NK-cell and T-cell populations (**Figure 4A**). Notably, amongst the *CD4*+ T-cells was a robust population of regulatory T-cells (*RTKN2*, *FOXP3*, *AC133644.2*, *CD4*, *IL2RA*, *TIGIT*, *CTLA4*, *FCRL3*, *LAIR2* and *IKZF2*) (**Figure 4B** and **4C**). This population represented over 10% (227/2126) of the detected *CD4*+ T-cells. Since the PBMC reference does not include exhausted T-cells, to detect this subpopulation, we performed subclustering of the *CD8*+ effector memory T-cells (CD8TEM), and looked at the expression of traditional exhaustion markers, specifically *PDCD1*, *LAG3*, and *TOX* (**Figure 4D** and **4E**). We found that the largest subpopulation of CD8TEM cells had higher expression of all three marker genes, indicating the robust presence of the exhausted T-cell population. Given that CD8TEM also represents the majority of *CD8*+ T-cells within the OS samples, T-cell exhaustion within osteosarcoma appears to be prevalent (1249/2610, 48%). Taken together, the significant population of regulatory and exhausted T-cells could be one of the major contributors to the immunosuppressive TME in OS.

### Osteosarcoma

Leveraging the large-scale copy number alterations that serve as the signature of osteosarcoma, we utilized *inferCNV* (37) to detect relevant subclones of OS cells with distinct copy number alterations. We observed several subpopulations with normal arm-level copy number profiles, indicating that normal fibroblast or mesenchymal cells likely existed within our osteosarcoma clusters. In order to remove these, we combined all six OS cell clusters potentially containing normal cells derived from the primary clustering analysis shown in **Figure 1**, and then subclustered the OS cells as a whole (**Supplementary Table S8**) prior to sample-specific *inferCNV* analysis. Subclustering results in 8 subclusters, including two subclusters of normal cells (subclusters 6 and 8) (**Supplementary Figure 2**). After removal of normal cell population, *inferCNV* analysis was carried out on individual patient basis. Between 3 and 8 subclones were observed in patients, with one sample having too few detected osteosarcoma cells to perform *inferCNV*. Several arm level copy number alterations were found recurrently in subclones of OS cells across samples, including CNVs previously reported in osteosarcoma. These CNVs include amplifications of chromosome arms 1p (38), 1q (38–42), 7p (43), 8q (38,41,44–49), and 20p (50–52), as well as deletions of 6q (53,54).

### Cell-cell interactions

Next, we sought to investigate how the various cell types identified in our scRNA-seq analysis interact within the OS TME to determine their possible role in immunosuppression. We utilized *CCCExplorer* (55) to examine ligand-receptor interactions using a curated database of known interacting pairs. We queried *CCCExplorer*’s database for interactions based on differentially expressed genes calculated using Seurat’s *FindMarkers* function (**Figure 5, Supplementary Table S9**). We looked specifically at signaling interactions involving from T-regs, OS, MDSCs, and macrophages in order to determine how this signaling may contribute to immunosuppression within the tumor, prioritizing interactions that also had downstream pathway expression support.

**Figure 5:**
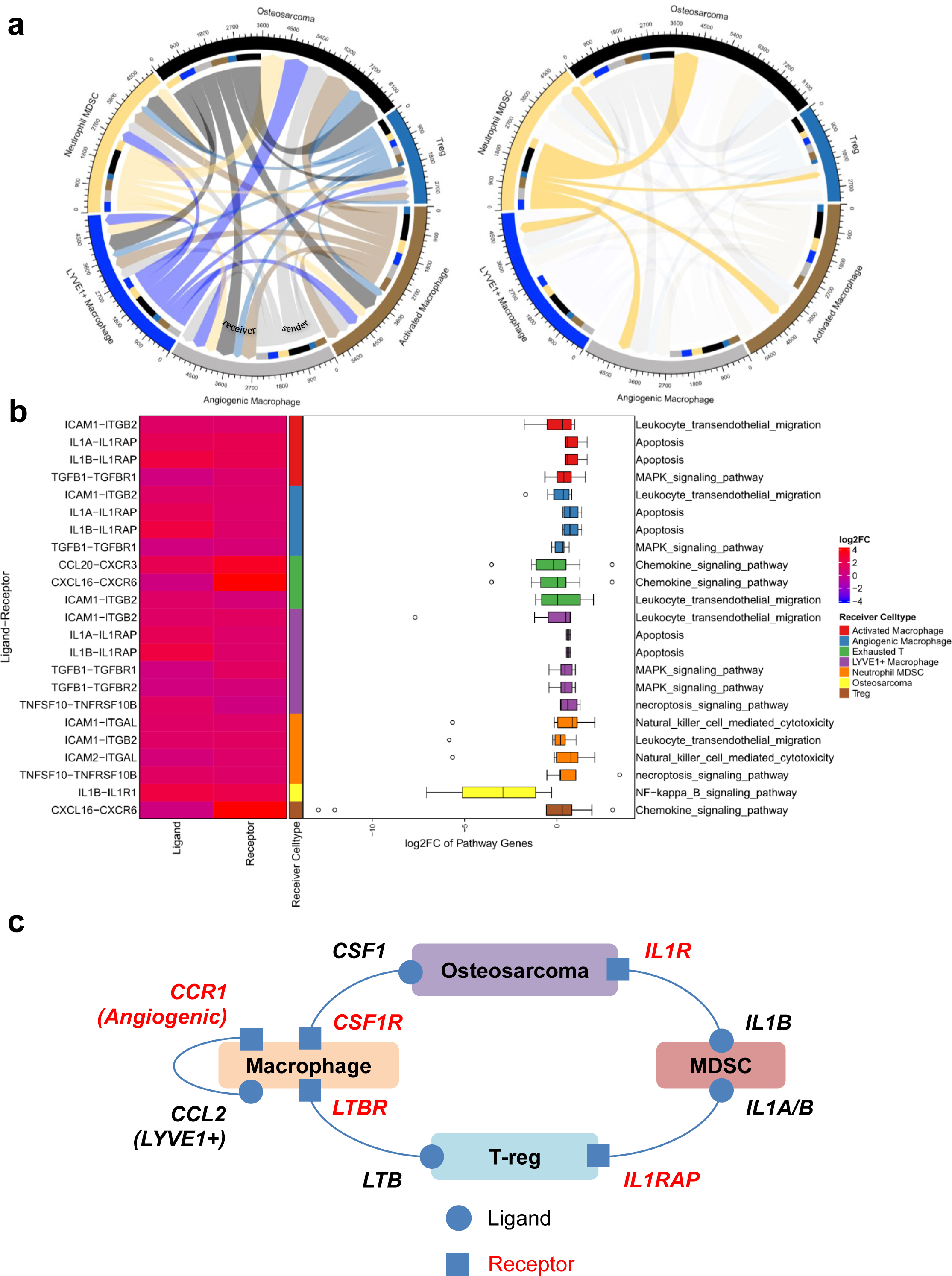
Upregulated ligand-receptor interactions from *S2C2*, separated by ligand-expressing cell type. **(A)** (left) A chord diagram from ligand-receptor interaction network between tested cell types. (right) A chord diagram highlighting the interaction originating from myeloid-derived suppressor cells (MDSC). Width of links are proportional to the number of ligand-receptor interactions identified. **(B)** Ligand-receptor interactions and downstream pathway expression originating from myeloid-derived suppressor cells (MDSC). Only identified ligand-receptor interactions with significant enrichment of the downstream pathway are shown. (left) Ligand and receptor expression, annotated by receiver cell type. (right) Boxplot showing expression of genes in pathways downstream of the ligand-receptor interaction in the receiver cell type. **(C)** Summary of selected interactions between immune cell types and osteosarcoma.

We found robust interaction between MDSCs and other cell types, including multiple interactions known to be involved in tumorigenesis and immunosuppression. For example, *IL1B* was highly expressed in MDSC, and its receptor *IL1R1* was highly expressed in osteosarcoma (**Figure 5B, C**). This interaction leads to downstream upregulation of *NF-κB* through *CHUK* upregulation, which is known to be highly expressed in osteosarcoma and is implicated in tumorigenesis, and whose inhibition lead to apoptosis and repressed proliferation and invasiveness in cell line models(56). MDSCs expressing *IL1A/B* were also predicted to interact with several macrophage subtypes (angiogenic, activated, LYVE1+) through *IL1RAP*. IL1RAP expression has been identified on tumor and stromal cells of multiple tumors (57), promoting an immunosuppressive microenvironment, and its interaction with IL1A/IL1B has also been specifically targeted in a phase 1b clinical trial (58).

Regulatory T-cell signaling to macrophages was observed. We identified *LTB* overexpression in regulatory T-cells, which interacts as a ligand with *LTBR*, which was highly expressed in activated and angiogenic macrophages (**Figure 5C, Supplementary Figure S5**). Ligand-receptor binding between the two activates the noncanonical NF-κB pathway, which also showed evidence of upregulation of downstream genes, and which affects inflammation in the tumor microenvironment of several cancers (59,60).

We also saw signaling between osteosarcoma cells and macrophage populations. OS cells expressed *CSF1*, which has been shown to be associated with protumor activity of tumor-associated macrophages in multiple sarcomas including OS, and is a regulator of the proliferation and survival (61,62). This interacts with *CSF1R*, highly expressed on activated and angiogenic macrophages, and activates several downstream pathways (**Figure 5C, Supplementary Figure S6**). CSF1 secretion by OS cells results in the polarization of bone-marrow-derived macrophages toward an M2 phenotype, and targeting this interaction with a CSF1R inhibitor suppressed OS xenograft growth and lung metastatic potential (62), and is being explored in multiple cancers (63).

We also looked at interactions between other immune cells in osteosarcoma. We detected an interaction between *LYVE1*+ macrophages and angiogenic macrophages involving *CCL2* and *CCR1* (**Figure 5C, Supplementary Figure S7**). LYVE1-expressing macrophages have been shown to have a role in maintaining a proangiogenic perivascular niche in cancer (64), and CCL2 is the primary chemokine responsible for recruitment of angiogenic macrophages (65). *AKT3* upregulation downstream of this interaction in angiogenic macrophages may lead to inflammatory and angiogenic responses as has been shown previously in cancer (66).

### Spatial transcriptomics

Like other solid tumors, osteosarcoma has a complex tissue structure and heterogenous cellular organization. To understand potential immunosuppression from a spatial perspective in the OS TME, we studied the cellular organization and architecture of the tissue using spatial transcriptomics analyses with the 10X Genomics Visium platform on two pre-treatment primary tumor biopsy tissue samples. After quality control filtering, normalization, and data integration using *Seurat* (36) and *Harmony* (67), we used Robust Cell Type Decomposition (RCTD) (68) to infer spot-level cell composition and compare overall cell type frequencies in the samples.

Similar to the scRNA-seq data, these samples primarily consisted of OS cells and immune cells, along with populations of endothelial cells and osteoclasts (**Figure 6A**). Immune cell abundance varied between the samples, with some subtypes being more prevalent in one sample than the other (**Figure 6B**). For instance, activated macrophages, angiogenic macrophages, and neutrophils/monocytes were more prevalent in Patient A, whereas CD8+ T-cells, CD4+ T-cells, and IFN-stimulated macrophages were observed more in Patient B.

**Figure 6:**
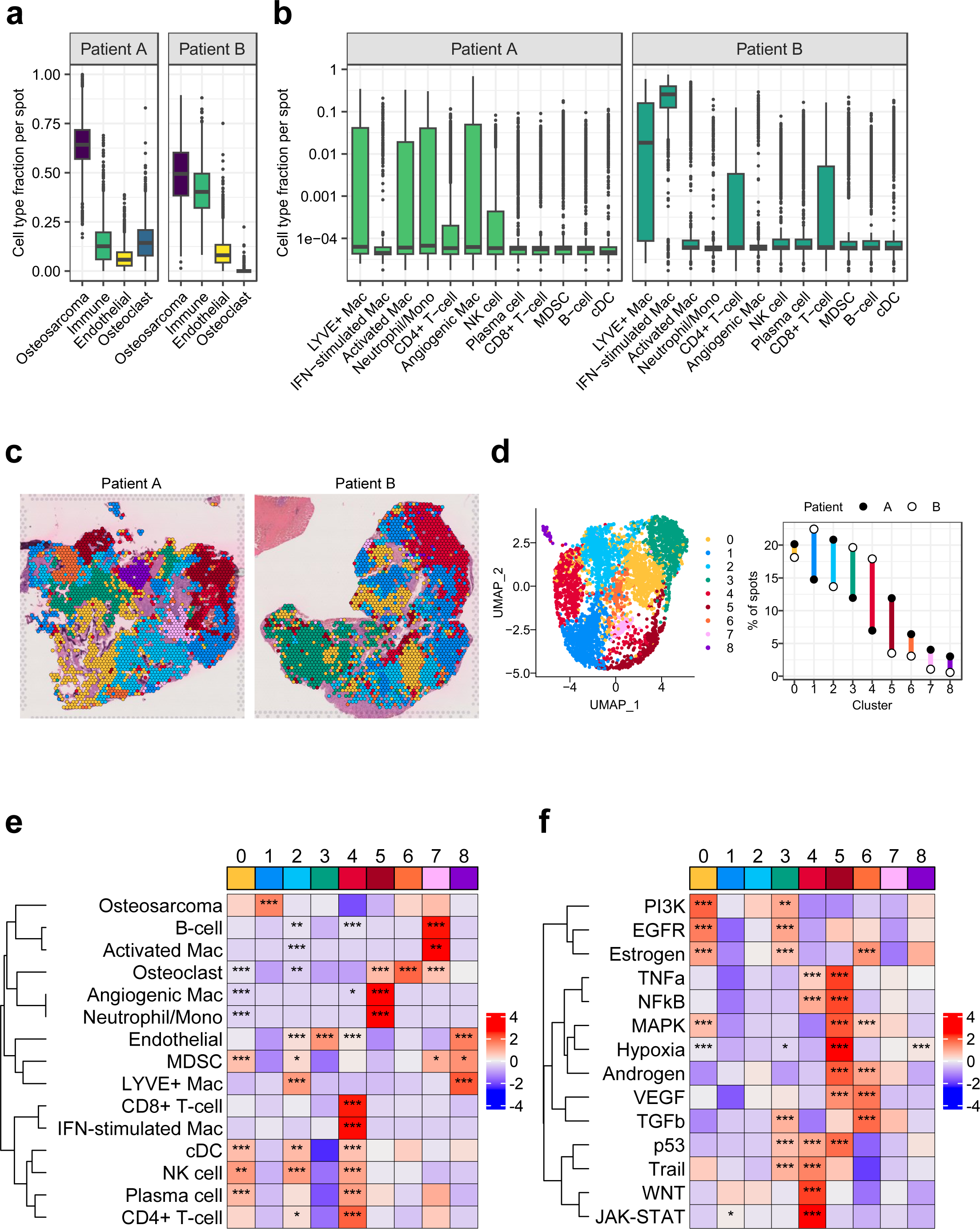
Spatial transcriptomics analysis of osteosarcoma **(A)** Estimated fractions of major classes of cell types per spot as determined by deconvolution using RCTD and the single cell reference dataset. The Immune class was aggregated from values inferred for 12 subtypes of immune cells. Osteosarcoma, Endothelial, and Osteoclast were calculated directly. **(B)** Estimated fractions of 13 immune cell subtypes per spot inferred by deconvolution showed variability between patients. **(C)** H&E images overlayed with spot cluster membership determined by unsupervised clustering of gene expression defining regional patterning. **(D)** Cluster membership information depicted as a UMAP plot (left) and as a percent of total spots for each patient (right). **(E)** Heatmap with z-scores based on the median cell type fraction per cluster. **(F)** Heatmap with z-scores based on the median PROGENy scores per cluster. These scores of relative pathway activity based on weighted gene expression levels were calculated per spot. Asterisks indicate the adjusted significance level for enrichment determined by one-sided Wilcoxon rank sum test. Spots in a single cluster were compared to all other spots. *, P<0.05; **, P<0.01; ***, P<0.001.

To better understand the cellular distribution of various cell types, eight clusters with unique gene expression patterns were identified by unsupervised clustering of spots (**Figure 6C and 6D**). Four clusters (0–3) defined the largest area of the tumors and were common to both samples, whereas other five clusters showed tumor-specific localizations (**Figure 6C and D**). Cell composition contributed to these regional identities based on the enrichment of estimated cell type fractions with each cluster (**Figure 6E**). Both tumors had comparable regions enriched in MDSCs (cluster 0), OS cells (with fewer immune cells, cluster 1), LYVE+ macrophages (cluster 2), or endothelial cells (cluster 3). Patient A had a region marked by neutrophils/monocytes and angiogenic macrophages (cluster 5) and another enriched in activated macrophages and B cells (cluster 7). In contrast, Patient B had a region enriched in CD8+ T-cells, CD4+ T-cells, and IFN-stimulated macrophages (cluster 4). These differences in immune cell composition between samples were largely attributable to localized phenomena.

To understand the biological significance and phenotypes of different regions within the samples, we ran signaling pathway activity analysis – *PROGENy*. Different clusters showed distinguished gene expression patterns associated with cancer-relevant signaling pathways (**Figure 6F**). Common to both tumors, relatively large areas showed gene expression patterns associated with EGFR (cluster 0) and TGFB (cluster 3) signaling pathway activity. Both pathways have been associated with immunosuppression in cancer and response to immunotherapy (69,70). Consistent with this, these areas contained immunosuppressive cells like MDSCs (cluster 0) and generally lacked enrichment of anti-tumor cells, like CD8+ T cells. In one sample, we observed a hypoxia-associated gene expression pattern (cluster 5), which also had its own distinct enrichment of immune cell types, including angiogenic macrophages and neutrophils/monocytes clustered with osteoclasts. Finally, the region with the clearest enrichment of CD8+ T cells (cluster 4) lacked enrichment of the EGFR, TGFB, and hypoxia-associated gene expression signatures. Overall, these data, for the first time, showed the spatial distribution of different cell types and cell states, particularly in macrophages, in OS TME. It also suggests that the TME of OS is largely immunosuppressive and its architecture consists of potentially distinct immunosuppressive milieus.

## Discussion

The suboptimal result of recent immunotherapy clinical trials suggested that OS has a complex immune TME. To improve the efficacy of treatments, particularly immunotherapy, a better understanding of the OS immune landscape is urgently needed. To address this, we performed scRNA-seq analysis of six treatment naïve OS samples and combined with the data with published scRNA-seq data of other six treatment naïve OS samples to generate the largest scRNA-seq dataset for OS to date. We aimed to better understand the potential immunosuppressive TME in OS by focusing on immune cells identified in OS TME.

With the combined dataset, we identified similar immune cell types reported in previous OS scRNA-seq studies (11,12), including immunosuppressive related cell types, e.g., CD4^+^ regulatory T cells and CD8^+^ exhausted T cells. These regulatory and exhausted T-cells play well-established roles in the microenvironment of multiple tumors, including osteosarcoma (11–13). T-cell exhaustion is a hallmark of immune evasion in many cancers, and overcoming this phenomenon in solid tumors is essential for the efficacy of emerging treatments such as CAR-T therapy (71,72).

Notably, although previous studies showed the presence of neutrophils in OS, we identified two subtypes of neutrophils including a major subtype expressing previous identified myeloid-derived suppressor cell (MDSC) markers. MDSCs contribute to the pathological immune response in cancer and may be potent targets for immunotherapy development (73). In addition, Zhou et al. (12) noted an increased presence of neutrophils in primary tumors as compared to recurrent tumors using a chemotherapy-treated cohort. Our treatment-naïve cohort of primary tumors also showed a robust population of neutrophils, and given that the majority were immunosuppressive, this likely represents an important axis of immune evasion particularly for primary osteosarcoma and warrants further investigation.

We also found a population of LAMP3+ cDCs within our cohort, consistent with previous findings in osteosarcoma. These cDCs interact with regulatory T cells to perform their immunosuppressive functions (13,23). Liu et al. (13) highlighted the interaction between LAMP3+ mregDCs and regulatory T-cells through the interaction of CD274 and PVR with PDCD1 and TIGIT, respectively., specifically within tumors. This underscores the importance of mregDCs in mitigating antitumor immunity in osteosarcoma. Our findings confirm the common presence of mregDCs in osteosarcoma, emphasizing the importance of further study of this immunosuppressive cell type.

The presence of immunosuppressive and angiogenic TAMs in osteosarcoma was also found in our dataset. Building on the previous analyses (11,12), we examined M1-M2 macrophage polarity amongst our macrophage subclusters using the markers from those studies, but found no evidence of MKI67+ tissue resident macrophages (**Supplementary Figure S4**). We did not observe strong evidence of M1 and M2 macrophage polarity in our dataset which aligns with the argument that macrophages have more complex phenotype than M1 and M2 polarity. In our analysis, distinct phenotypic macrophages were found, including classic activated macrophages, angiogenic macrophages, and interferon-stimulated macrophages, based on distinct markers for each subcluster. Additionally, subclustering revealed a distinct population of LYVE1+ angiogenic macrophages, previously found in several cancer subtypes (17–21). Although the exact role of each macrophage subtype in OS TME remain unclear, as the most abundant immune cell types with distinct phenotypes, macrophages are attractive targets for future immunotherapy development. Liu et al. previously (11) identified four major subpopulations of osteoclast: mature osteoclasts (*CTSK*, *ACP5*, *MMP9*), proliferative progenitor osteoclasts (*CD74*, *CD14*, *HLA-DRA*, *MKI67*, *CTSK^lower^*, *ACP5^lower^*, *MMP9^lower^*), and non/hypofunctional osteoclasts with low expression of such markers. While not a focus of this study, our subclustering of osteoclasts revealed a population of progenitor osteoclasts positive for progenitor markers *CD74*, *CD14*, *HLA-DRA* and *MKI67*, as well as the proliferation marker *TOP2A*, matching the previous results (**Supplementary Figure S3**). The other four populations of osteoclasts seemed to closely match the more mature osteoclasts subtypes with few distinctive differentially expressed genes, and there were no strong indications of separate hypofunctional or nonfunctional osteoclast subtypes. Further analysis of the mature osteoclast subtypes may reveal unique phenotypes.

Using CCCExplorer and scRNA-seq data from various cell types, we identified robust interactions between MDSCs and other cells, including macrophages through IL1RAP, leading to NF-κB upregulation and an immunosuppressive microenvironment. Regulatory T-cells signaled to macrophages via LTB and LTBR, activating the noncanonical NF-κB pathway. Osteosarcoma cells polarized macrophages to an M2 phenotype via CSF1 and CSF1R, and interactions between LYVE1+ and angiogenic macrophages through CCL2 and CCR1 promoted a proangiogenic niche and inflammatory responses.

We further examine spatial transcriptomics on two pre-treatment biopsy samples to verify the presence of these immunosuppressive cell types and examine how they colocalize within osteosarcoma. We found distinctions between those samples in their immune cell populations, which broadly indicate variations in interferon stimulation and macrophage subtypes. Osteosarcoma cells, MDSCs, and *LYVE1*+ macrophages were comparable between the two samples, indicating commonalities between tumors. However, the importance of the different immune cell population frequencies between these two tumors, which also showed differences in pathway expression related to growth factor expression, highlight the need for more spatial transcriptomics data to explore cellular heterogeneity in osteosarcoma and its impact on treatment and survival outcomes.

Overall, our findings suggest a broad network of pathological immunosuppression within osteosarcoma. While further orthogonal validation and functional studies are necessary to confirm these findings, it would be prudent to explore treatment strategies that address the observed immunosuppressive signaling within osteosarcoma.

## Material and Methods

### Sample Collection and Tissue Dissociation

Tumor tissue specimens of six OS patients were collected at Connecticut Children’s Medical Center (CCMC) or Children’s Hospital Colorado (CHCO). All specimen collection and experiments were reviewed and approved by the Institutional Review Board of CCMC or CHCO, respectively. Written informed consent was obtained prior to acquisition of tissue from patients. Diagnosis was confirmed by pathologist assessment. The specimens were stored in MACS tissue storage solution (Miltenyi Biotec, Bergisch Gladbach, Germany). The three CCMC samples were transferred to The Jackson Laboratory for Genomic Medicine for the tissue dissociation process.

For CCMC samples, the tissue specimens were minced and enzymatically dissociated in DMEM medium supplemented with Collagenase II (250U/ml, Gibco #17101-015, Waltham, Massachusetts, USA) and DNase (1μg/ml, Stemcell #07900, Cambridge, Massachusetts, USA) for up to 3 cycles of 37 °C 15 minutes dissociation with agitation. After each cycle, undigested tissue pieces were settled down by gravity and supernatant was transferred to cold Buffer I solution (10% FBS/DMEM medium supplemented by EDTA (2mM) and 2% BSA (Lampire #7500854, Pipersville, Pennsylvania, USA). Fresh dissociation solution was applied to the undigested tissue for next digestion cycle. Cells collected from each cycle were merged, spun down and resuspended with ACK lysis buffer (Gibco #A1049201) to remove red blood cells. After 3 minutes on ice, lysis reaction was quenched by adding Buffer I solution. After centrifugation, cells were resuspended in Buffer I solution and strained through a 70-μm cell strainer. Cells were stained with propidium iodide (PI) and Calcein Violet (Thermo Fisehr Scientific). PI-negative and Calcien Violet-positive viable cells were sorted out using FACSAria Fusion system (BD Biosciences, Franklin Lakes, New Jersey, USA).

For CHCO samples, tissue specimens were viably cryopreserved in 90% FBS 10%DMSO immediately after collection. Tissue specimens were thawed at 37 °C, then enzymatically dissociated in a 0.1% DNase and Liberase 400μg/mL (Roche/Sigma-Aldrich, St. Louis, Missouri, USA) cocktail. Cells were then selected for viability using the FACSAria I cell sorter (BD Biosciences) in the Allergy and Clinical Immunology Flow Cytometry Facility at the Division of Allergy and Clinical Immunology, University of Colorado School of Medicine.

### Single-Cell Capture, Library Preparation and RNA-seq

Cells were washed and resuspended in PBS containing 0.04% BSA. Cells were counted on Countess II automated cell counter (Thermo Fisher, Waltham, Massachusetts, USA), and up to 12,000 cells were loaded per lane on 10x Chromium microfluidic chips (10x Genomics, Pleasanton, California, USA). Single-cell capture, barcoding, and library preparation were performed using the 10x Chromium X version 3 chemistry (74), and according to the manufacturer’s protocol. cDNA and libraries were checked for quality on Agilent 4200 Tapestation and quantified by KAPA qPCR before sequencing using a Novaseq 6000 (Illumina, San Diego, California, USA) v1.5 cycle flow cell lane at 100,000 reads per cell, with a 28-10-10-90 asymmetric read configuration.

### Single-Cell Data Processing and Quality Control

10X Genomics *Cell Ranger* (3.0.2) was used for read alignment (version 3.0.0, GRCh37) and count matrix generation (https://www.10xgenomics.com/support/software/cell-ranger/latest). Reads from each sample were initially filtered using multiple criteria: during FASTQ generation, reads with more than one mismatch in the 8bp i7 index are excluded. Only reads aligned to annotated transcripts with MAPQ scores greater than 255 are retained. Reads containing bases with Q30 scores below 3 are also excluded.

Following alignment, cell barcodes were filtered against a whitelist of barcodes provided by 10X Genomics (<2 mismatches). Barcodes associated with cells are distinguished from those associated with ambient mRNA using an adaptively computed UMI threshold via *Cell Ranger*, and a digital counts matrix is generated for each sample. Single-cell RNA expression data from an additional 6 samples described in Liu et al. (11) was downloaded from the Gene Expression Omnibus (GSE162454).

For all samples, additional preprocessing was performed using *Seurat* (version 4.0.5) (36) in R (version 4.1.1) (75). For each cell, the percentage of reads mapping to mitochondrial and ribosomal genes was calculated, and cells were filtered out according to the following criterion: >20% mtRNA, >50% rRNA, or <500 features expressed. Doublet detection and removal was performed using the *R*-package *DoubletFinder* (default parameters) (76). Read counts for each cell were log-normalized (scale factor=1e6), and the top 2000 variable features (selection method=“vst”) in the data set were calculated. Data was scaled and corrected for cell cycle score and percentage of mitochondrial RNA (“CellCycleScoring” and “ScaleData” functions, vars.to.regress = c(“percent.mt”, “S.Score”, “G2M.Score”)) prior to principal component analysis (PCA). *Harmony* (version 0.1.0) (67) was used to correct the top 50 principal components for patient sex and sample source site prior to downstream clustering analyses.

### Clustering and Cell Type Determination

Primary cell clustering was performed using *Seurat*’s “FindClusters” function (*Harmony*-correct PCs = 40, res = 0.25). The “FindAllMarkers” function (only.pos=TRUE, min.pct = 0.25, logfc.threshold = 0.25, test.use = “bimod” (77)) was utilized to detect markers for each cluster, with major cell types annotated using a known set of genes, specifically Macrophage/DC (*MSR1*, *C1QC*, *FOLR2*), Osteosarcoma (*SATB2*, *IBSL*, *ALPL*), NK/T-cell (*CD3D*, *TRBC1*, *NKG7*), Osteoclast (*ACP5*, *CTSK*, *MMP9*), Monocyte/Neutrophil (*S100A8*, *S100A9*, *FCN1*), Fibroblast (*TAGLN*, *ACTA2*, *FAP*), Endothelial (*CLEC14A*, *PLVAP*, *VWF*), B-cell (*MS4A1*, *CD79A*, *BANK1*), Plasma cell (*IGHG1*, *IGLC2*, *IGHG4*), and Mast cell (*TPSB2*, *TPSAB1*, *CPA3*), as well as proliferation markers (*MKI67*, *TOP2A*, *TYMS*).

Subsequently, major cell types were further subclustered to detect cell subtypes. We used the “subset” function for major cell clusters, reperformed normalization (“CellCycleScoring” and “ScaleData” functions, vars.to.regress = c(“percent.mt”, “CC.Difference”)), and *Harmony* correction of PCs. Subclustered cell types include macrophage/DCs (PCs=30, res=0.2), osteosarcoma/fibroblasts (PCs=40, res=0.15), osteoclasts (PCs=30, res=0.1), neutrophils (PCs=30, res=0.1), and proliferative macrophages/DCs (PCs=30, res=0.3). For the NK/T-cell cluster, subtypes were identified using *Azimuth* (36). Cells within the NK/T-cell cluster were mapped to NK/T-cell L1 annotations within the reference proliferating blood mononuclear cell (PBMC) dataset (35), and cells with a confidence score >0.75 were retained for downstream analysis.

### Copy Number Analysis

In order to characterize overall copy number changes between the identified osteosarcoma populations, as well as to detect subclones within each major cluster, *inferCNV* (version 1.10.0) (37) as implemented on a sample-specific basis (cutoff=0.1, HMM=T, denoise=T, analysis_mode=“subclusters”, hclust_method=“ward.D2”, cluster_by_groups=TRUE). All normal cell populations were used for comparison.

### Cell-cell interactions

*CCCExplorer* (55) was employed for the comprehensive investigation of cell-cell interactions and their impacts on downstream pathways. The integration of a manually curated ligand-receptor database, transcription factor-target database, experimentally validated pathway database, and crosstalk algorithms facilitated a thorough examination of inter-cellular crosstalk. Gene expression changes in ligand-receptor pairs across selected cell types of interest, including various macrophages, T-cells, NK cells, dendritic cells, fibroblasts, and neutrophils, were explored using the CCCExplorer ligand-receptor database. The potential downstream effects on cell functions were assessed by applying a crosstalk algorithm to networks generated from KEGG, Ingenuity Pathway Analysis (IPA), and a transcription factor-target interaction database. Attention was given to pairs where elevated expression of ligands from sender cells and receptors from recipient cells was detected. Further analysis of these interactions involved the identification of significant pathway enrichments using permutation test (p < 0.05).

### Visium Spatial Gene Expression Analysis

Formalin-fixed, paraffin-embedded (FFPE) samples of two primary OS pre-treatment biopsy tissue samples (Patients A and B) were obtained from CCMC and stored at 4 °C in the dark. Prior to Visium transcriptomics, RNA quality of FFPE samples was determined by DV200 score using Agilent (Santa Clara, California) TapeStation 4200 High Sensitivity DNA ScreenTape. Tissue blocks with DV200 scores above 50% were used for downstream processing. Briefly, FFPE sections were placed on a 10x Visium FFPE Gene Expression slide, deparaffinized, H&E stained, then imaged in brightfield using a NanoZoomer SQ (Hamamatsu Photonics, Shizuoka, Japan) slide scanner, followed by incubation with human-specific probe sets provided by the manufacturer for subsequent mRNA labeling and library generation per the manufacturer’s protocol (10x Genomics, CG000407). Library concentration was quantified using a Tapestation High Sensitivity DNA ScreenTape (Agilent) and fluorometry (Thermofisher Qubit) and verified via KAPA qPCR. Libraries were pooled for sequencing on an Illumina NovaSeq 6000 200-cycle S4 flow cell using a 28-10-10-90 read configuration, targeting 100,000 read pairs per spot covered by tissue.

Illumina base call files for all libraries were converted to FASTQs using bcl2fastq v2.20.0.422 (Illumina). Whole Visium slide images were uploaded to a local OMERO server. For each capture area of the Visium slide, a rectangular region of interest (ROI) containing just the capture area was drawn on the whole slide image via OMERO.web, and OMETIFF images of each ROI were programmatically generated using the OMERO Python API. FASTQ files and associated OMETIFF corresponding to each capture area were aligned to the GRCh38-specific filtered probe set (10x Genomics Human Probeset v1.0.0) using the version 2.1.0 *Space Ranger* count pipeline (10x Genomics).

*Seurat* (version 4.3.0) (36) was used for normalization, filtering, dimensional reduction, and visualization of the resulting gene expression data. Each section was processed independently. Spots with fewer than 1500 UMIs and 500 genes were removed. Normalization was performed using the “SCTransform” function in *Seurat*, with batch correction performed using *Harmony* (version 0.1.1) (67). Unsupervised clustering was performed in *Seurat* using Leiden clustering (“FindClusters”) (78). Cell type fractions within spots were estimated using Robust Cell Type Decomposition (RCTD, *spacexr* package version 2.1) (68) and the single cell RNA-seq data as the reference. Sixteen cell types identified above were used, including three major cell classes (Osteosarcoma, Endothelial, and Osteoclast) and 13 immune cell subtypes. The “Immune” fraction shown was the sum of all the individual subtype contributions. Differences in cell type composition between clusters were determined via a one-sided Wilcoxon rank sum test (79) for positive enrichment. Differences in pathway activity were calculated using *PROGENy* (version 1.14) (80).

## Supporting information

Supplementary Table S1 - Sample characteristics.xlsx

Supplementary Table S2 - Frequency tables of cell types.xlsx

Supplementary Table S3 - Differentially expressed genes by major cluster.xlsx

Supplementary Table S4 - Macrophage subclusters.xlsx

Supplementary Table S5 - Proliferative macrophage, DC subclusters.xlsx

Supplementary Table S6 - Monocyte, Neutrophil subclusters.xlsx

Supplementary Table S7 - NK, T cell subtypes.xlsx

Supplementary Table S8 - Osteosarcoma, fibroblast subclusters.xlsx

Supplementary Table S9 - CCCExplorer ligand-receptor interactions.xlsx

Supplementary Figure S1 - Cell Proportions by Patient.pptx

Supplementary Figure S2 - inferCNV.pptx

Supplementary Figure S3 - Osteoclast Subtype Markers.pptx

Supplementary Figure S4 - Macrophage Subtype Markers.pptx

Supplementary Figure S5 - Regulatory T-cell Ligand Interactions.pdf

Supplementary Figure S6 - Osteosarcoma Ligand Interactions.pdf

Supplementary Figure S7 - LYVE1+ Macrophage Ligand Interactions.pdf

## Acknowledgements

We gratefully acknowledge the contribution of the Single Cell Biology service, the Genome Technologies service, the Histology Services, the Flow Cytometry Service, and cyberinfrastructure high performance computing resources at The Jackson Laboratory for expert assistance with the work described herein.

## Materials and Correspondence Statement

Correspondence should be addressed to Dr. Ching Lau and Dr. Stephen T.C. Wong.

## Data Availability

Internal single-cell RNA expression and Visium spatial transcriptomics data will be uploaded to the Gene Expression Omnibus upon final publication. Single-cell RNA expression data from the external cohort of 6 samples was downloaded from the Gene Expression Omnibus (GSE162454). Correspondence should be directed to Dr. Ching Lau and Dr. Stephen T. Wong.

## Supplementary Figures

*Supplementary Figure S1*: Major cell type proportions of each sample.

*Supplementary Figure S2*: *inferCNV* plots for each sample.

*Supplementary Figure S3*: Osteoclast subtype markers used by Liu et al. (11) plotted across our identified osteoclast subclusters.

*Supplementary Figure S4*: M1/M2/tissue-resident macrophage (TRM) subtype markers used by Liu et al. (11) and Zhou et al. (12) and plotted across our identified osteoclast subclusters.

*Supplementary Figure S5*: Ligand-receptor interactions and downstream pathway expression originating from regulatory T-cells cells.

*Supplementary Figure S6*: Ligand-receptor interactions and downstream pathway expression originating from osteosarcoma cells.

*Supplementary Figure S7*: Ligand-receptor interactions and downstream pathway expression originating from *LYVE1*+ macrophage cells.

## Supplementary Tables

*Supplementary Table S1*: Patient and sample characteristics.

*Supplementary Table S2*: Frequency table of the contribution to the primary clusters by (**a**) sample, (**b**) patient sex, and (**c**) cohort. (**d**) Major cluster membership of each cell.

*Supplementary Table S3*: Full gene list showing markers for each primary cluster.

*Supplementary Table S4*: (**a**) Frequency table of the contribution to the macrophage subclusters by sample. (**b**) Macrophage subcluster membership of each cell. (**c**) Cell markers of each macrophage subcluster.

*Supplementary Table S5*: (**a**) Frequency table of the contribution to the proliferative macrophage/DC subclusters by sample. (**b**) Proliferative macrophage/DC subcluster membership of each cell. (**c**) Cell markers of each proliferative macrophage/DC subcluster.

*Supplementary Table S6*: (**a**) Frequency table of the contribution to the monocyte/neutrophil subclusters by sample. (**b**) Monocyte/neutrophil subcluster membership of each cell. (**c**) Cell markers of each monocyte/neutrophil subcluster.

*Supplementary Table S7*: (**a**) *Azimuth* output showing the predicted cell types/subtypes of the NK/T cluster cells after mapping to the PBMC data. Cells that mapped to NK or T-cell clusters by Azimuth with a confidence score > 0.75 were retained for downstream analysis (**b**) Frequency table of the contribution to the NK/T-cell L2 subtypes by sample.

*Supplementary Table S8*: (**a**) Frequency table of the contribution to the osteosarcoma/normal fibroblast subclusters by sample. (**b**) Osteosarcoma/normal fibroblast subcluster membership of each cell. (**c**) Cell markers of each osteosarcoma/normal fibroblast subcluster.

*Supplementary Table S9*: *CCCExplorer* output table, showing significant ligand/receptor interactions between selected cell types.

