## Supplementary figures and images for "Immunosuppressive tumor microenvironment of osteosarcoma"

### Supplementary Figure S1 - Cell Proportions by Patient.pptx

#### Slide 1
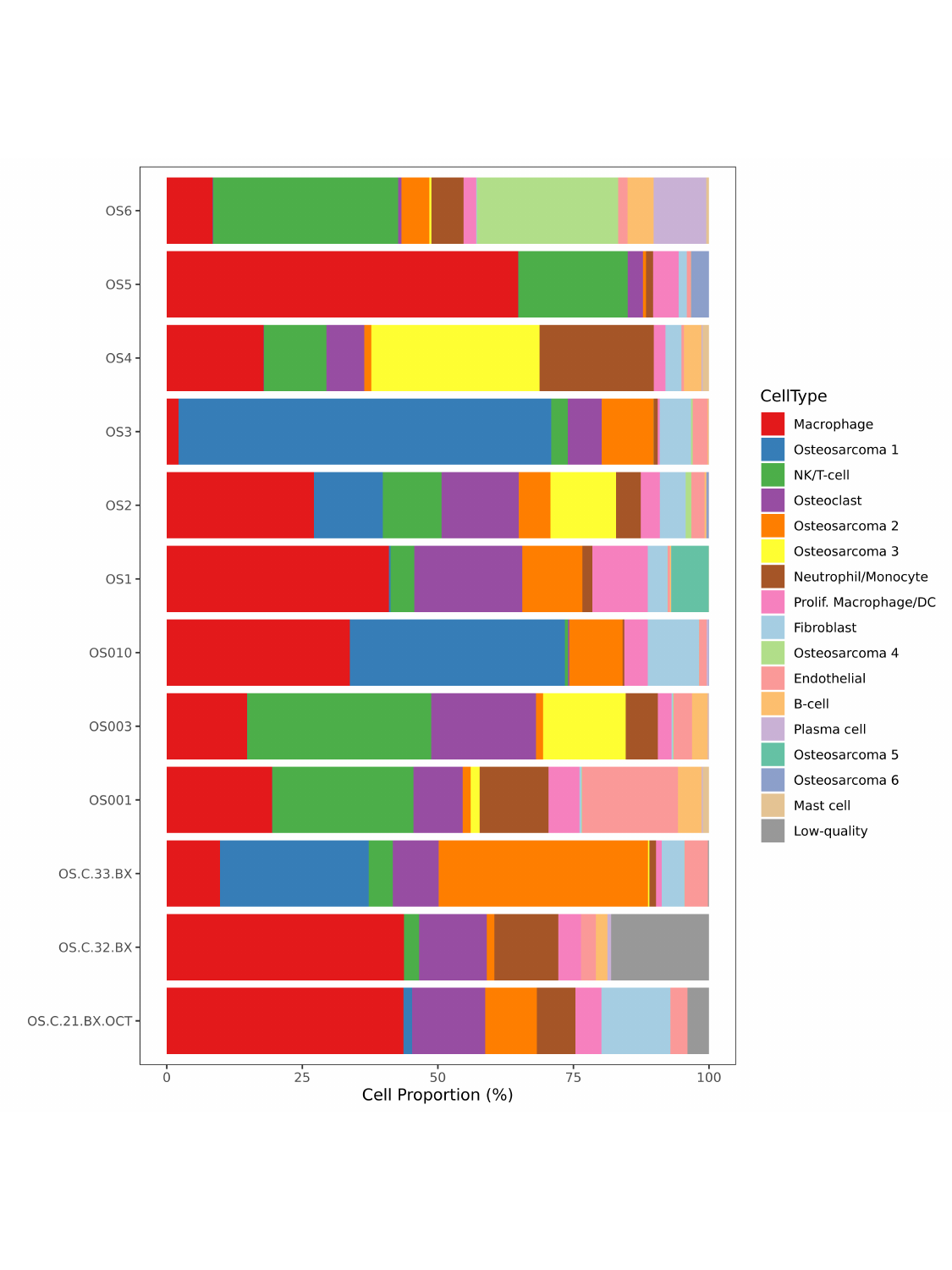

### Supplementary Figure S3 - Osteoclast Subtype Markers.pptx

#### Slide 1
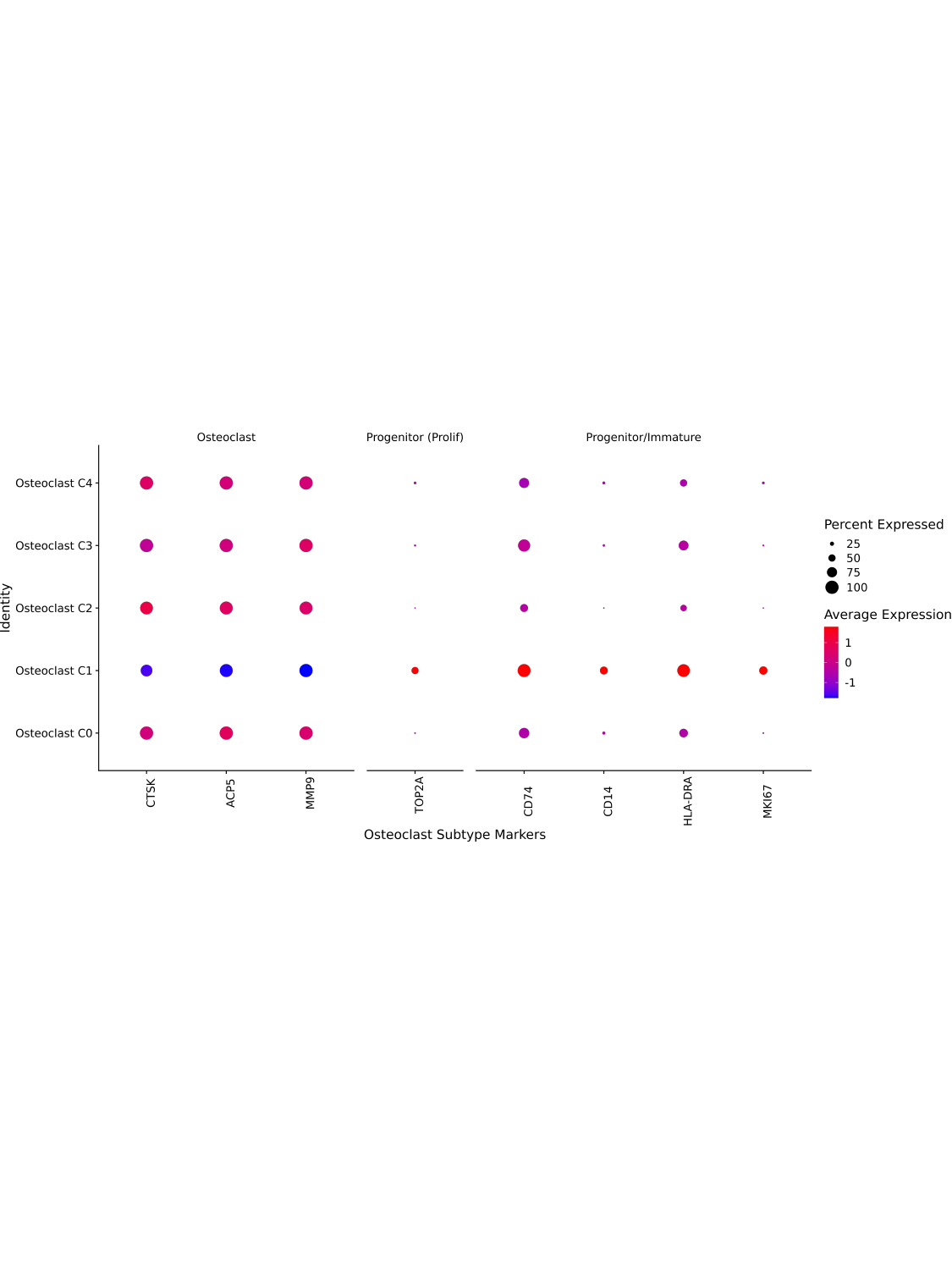

### Supplementary Figure S4 - Macrophage Subtype Markers.pptx

#### Slide 1
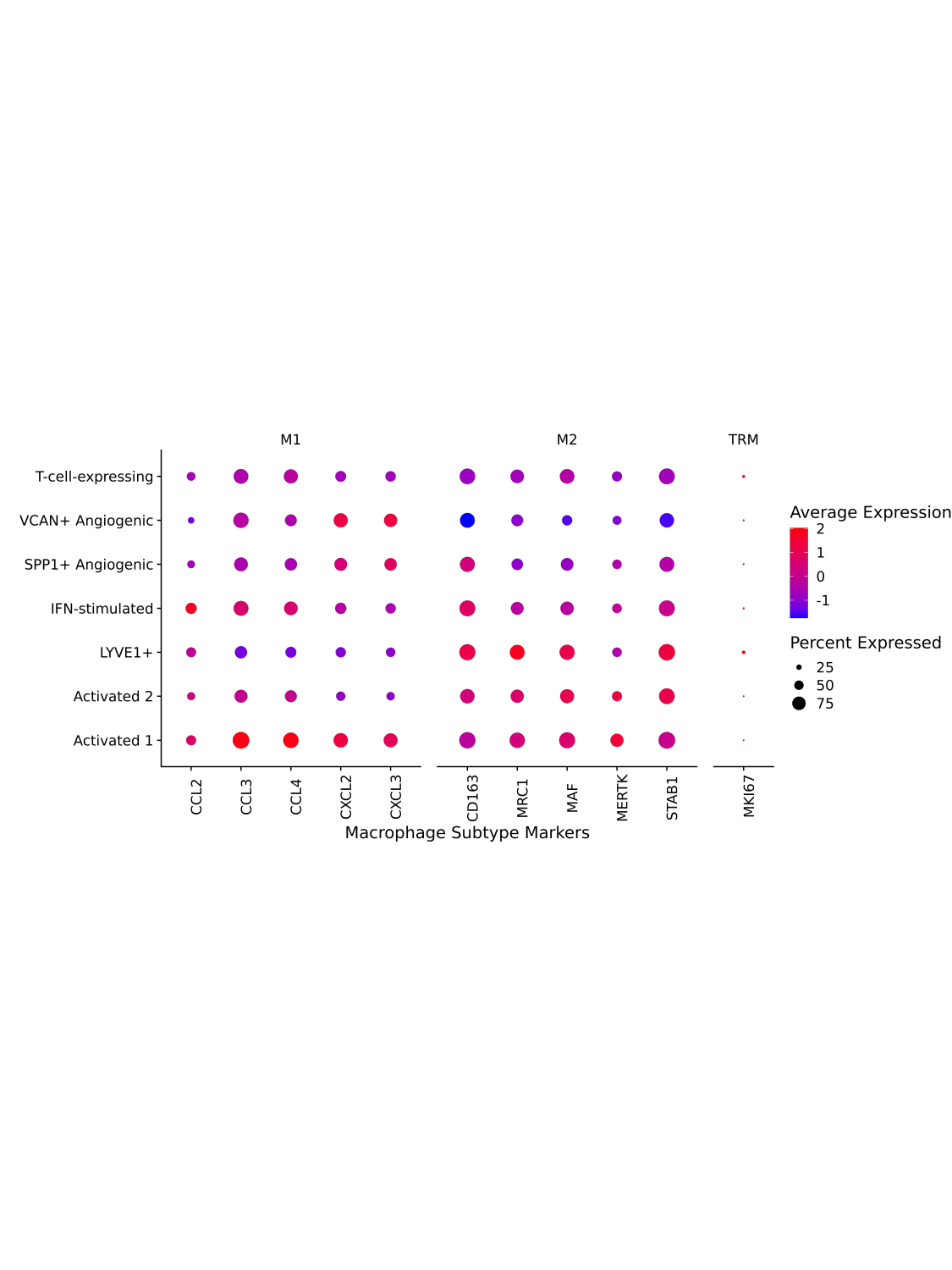

### Supplementary Figure S5 - Regulatory T-cell Ligand Interactions.pdf

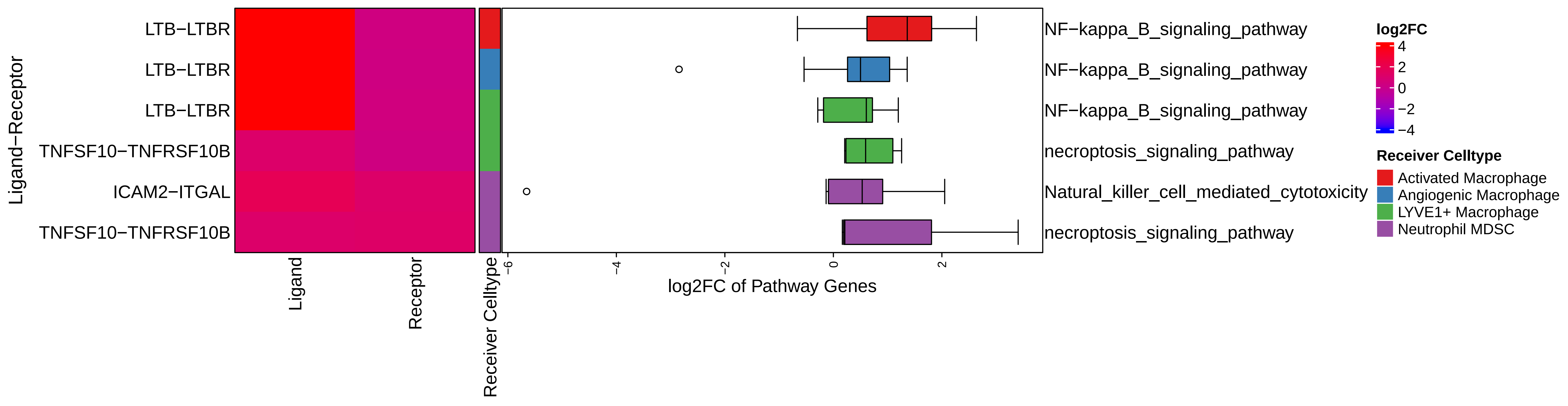

### Supplementary Figure S6 - Osteosarcoma Ligand Interactions.pdf

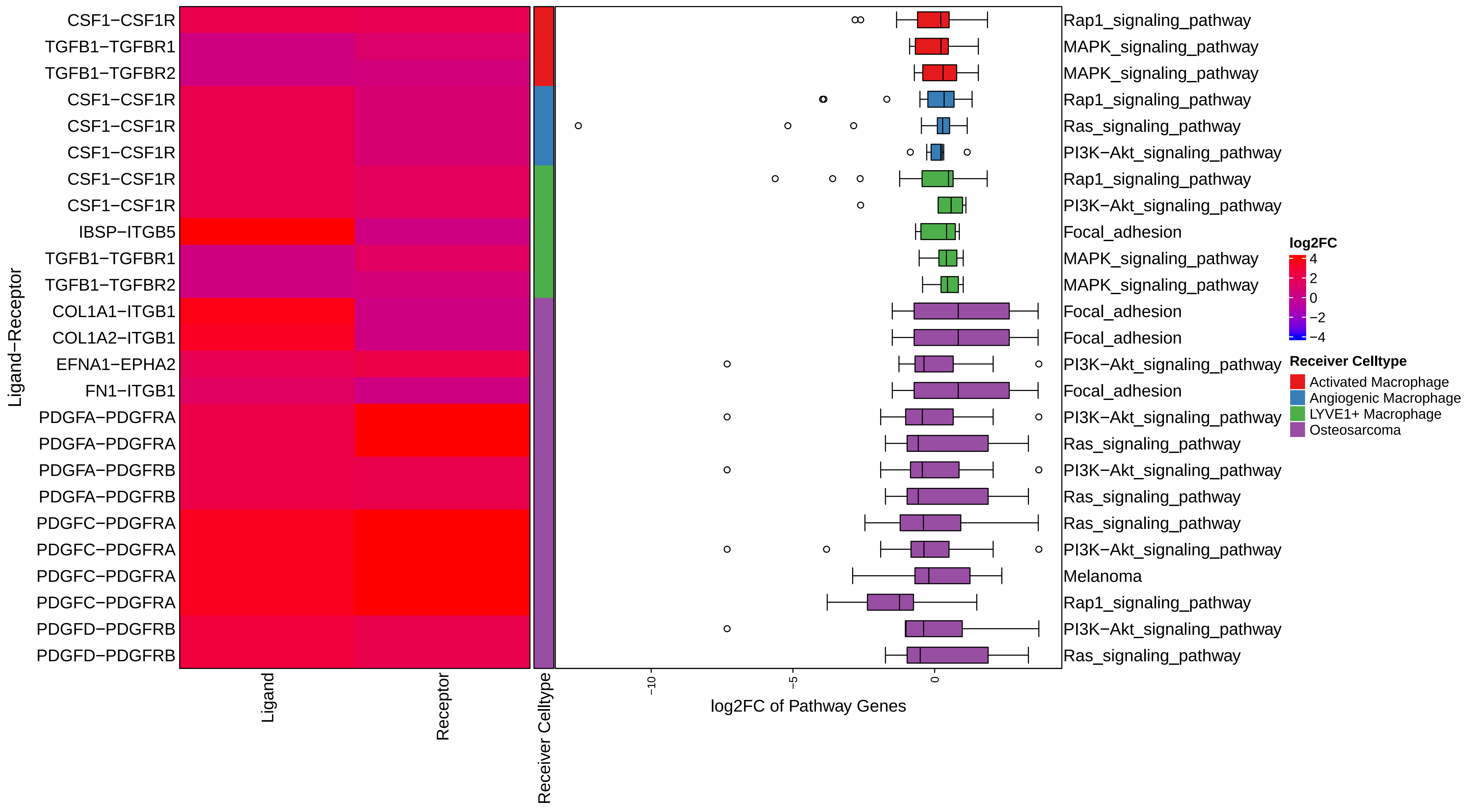

### Supplementary Figure S7 - LYVE1+ Macrophage Ligand Interactions.pdf

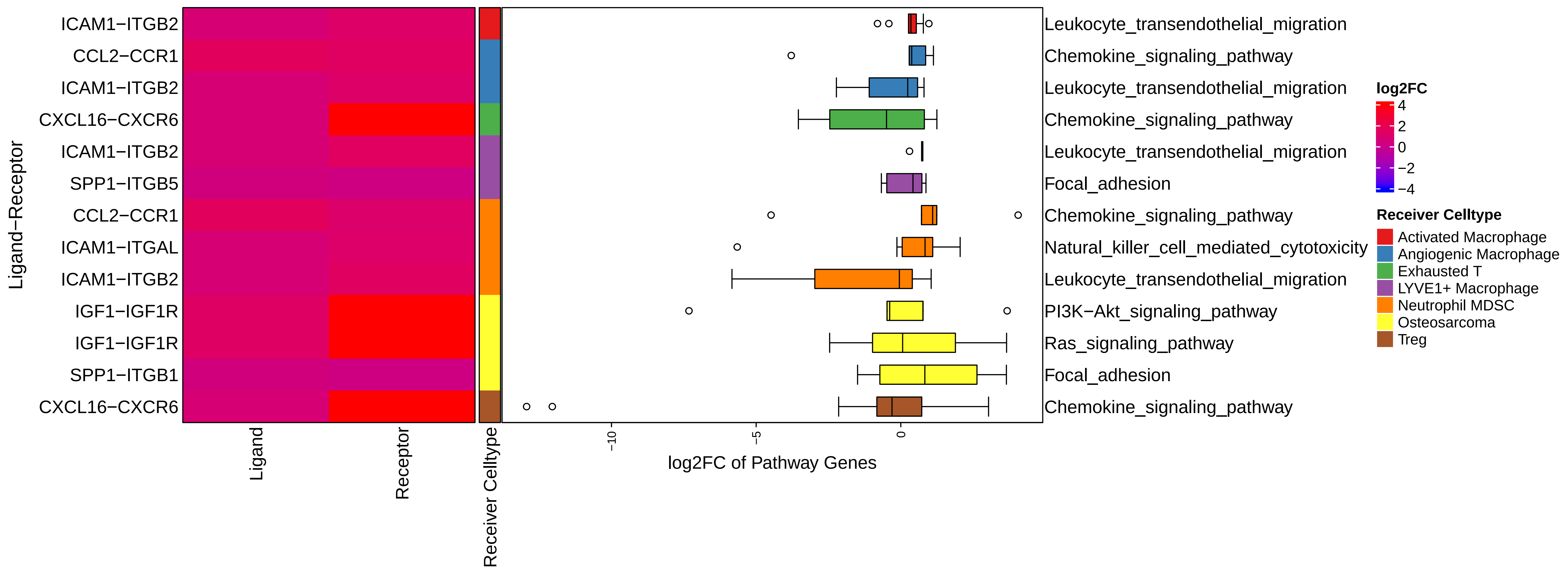
