## Supplementary Figure S2 - inferCNV.pptx for "Immunosuppressive tumor microenvironment of osteosarcoma"

### Slide 1
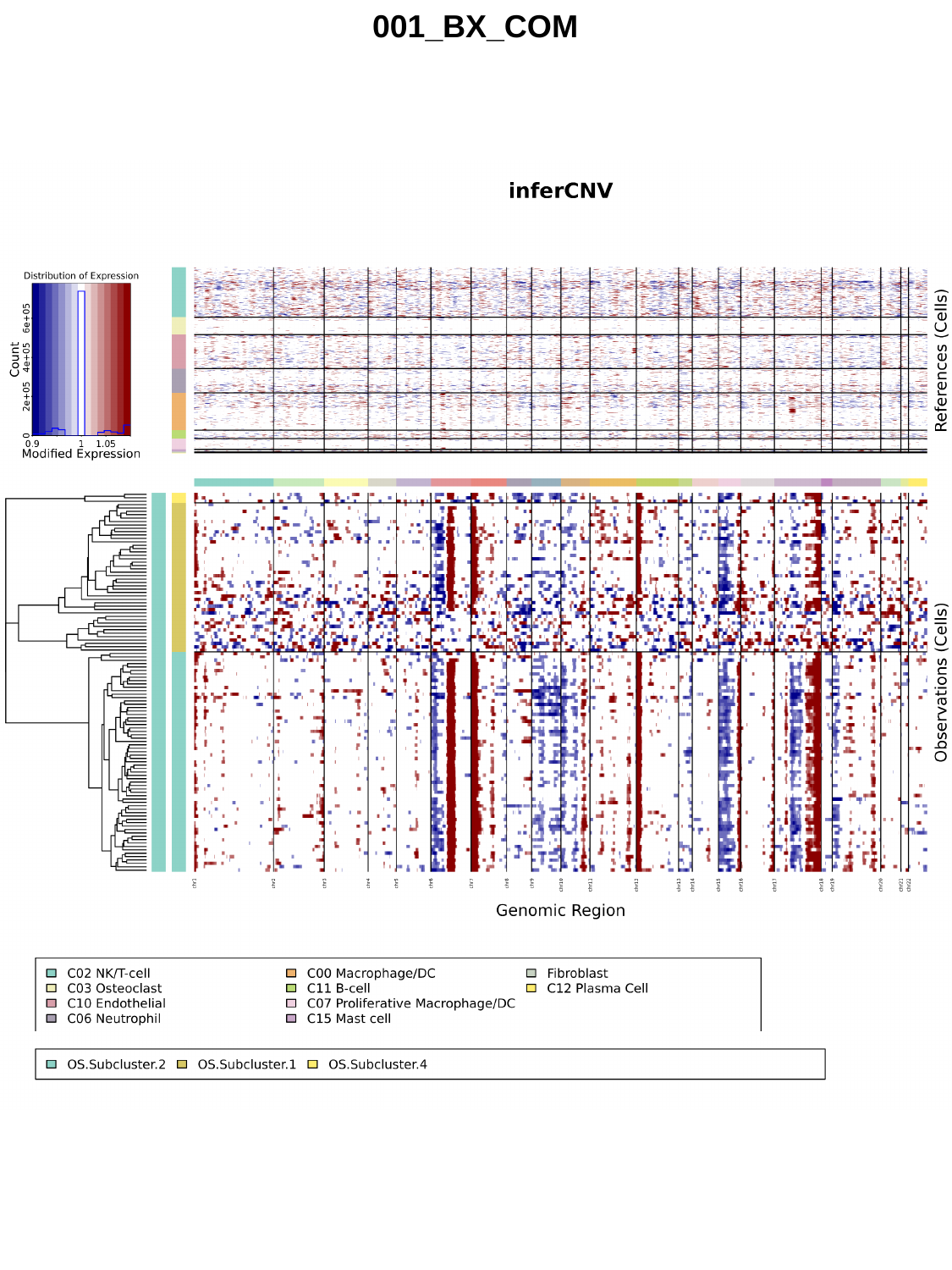

001_BX_COM

### Slide 2
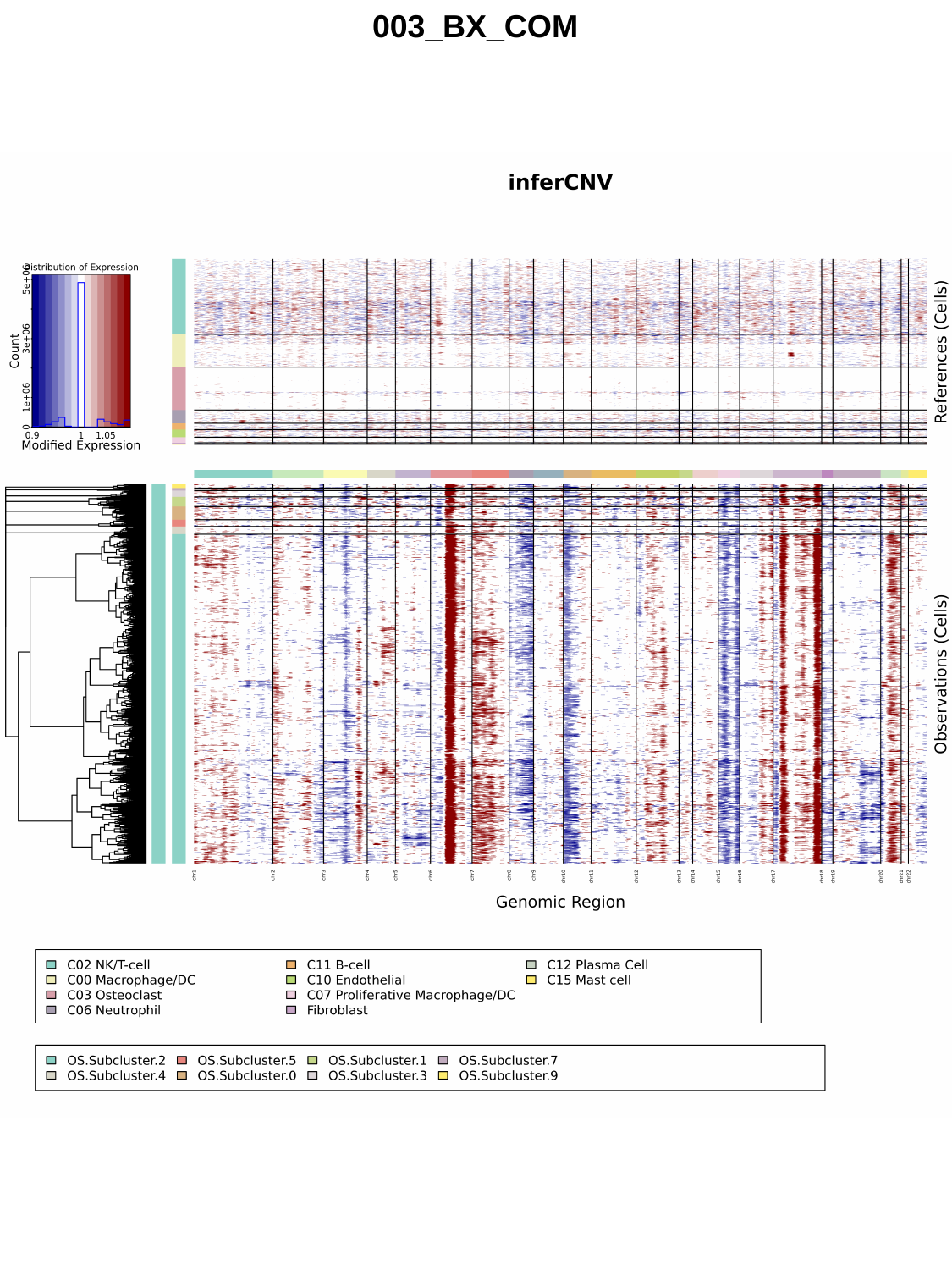

003_BX_COM

### Slide 3
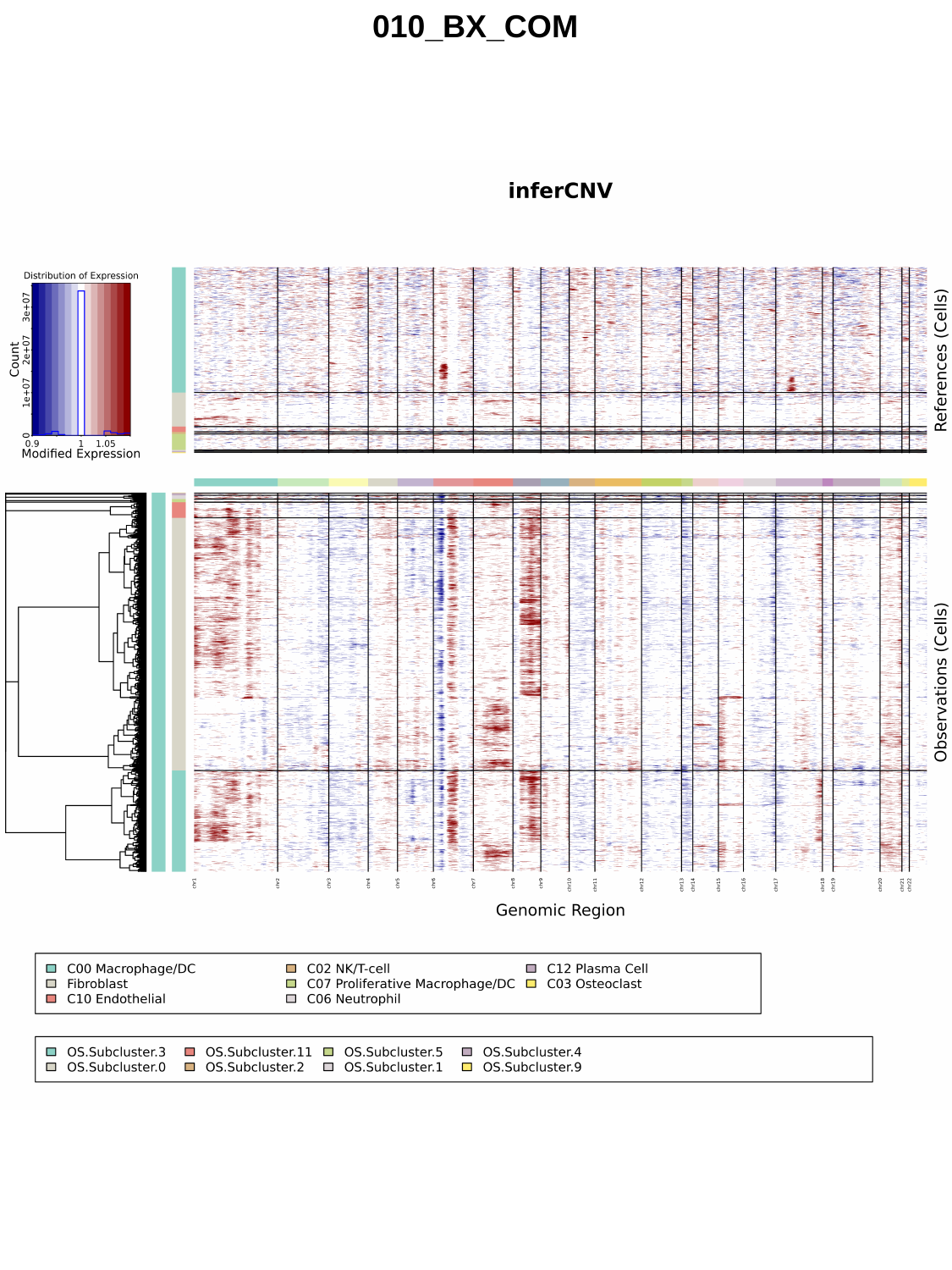

010_BX_COM

### Slide 4
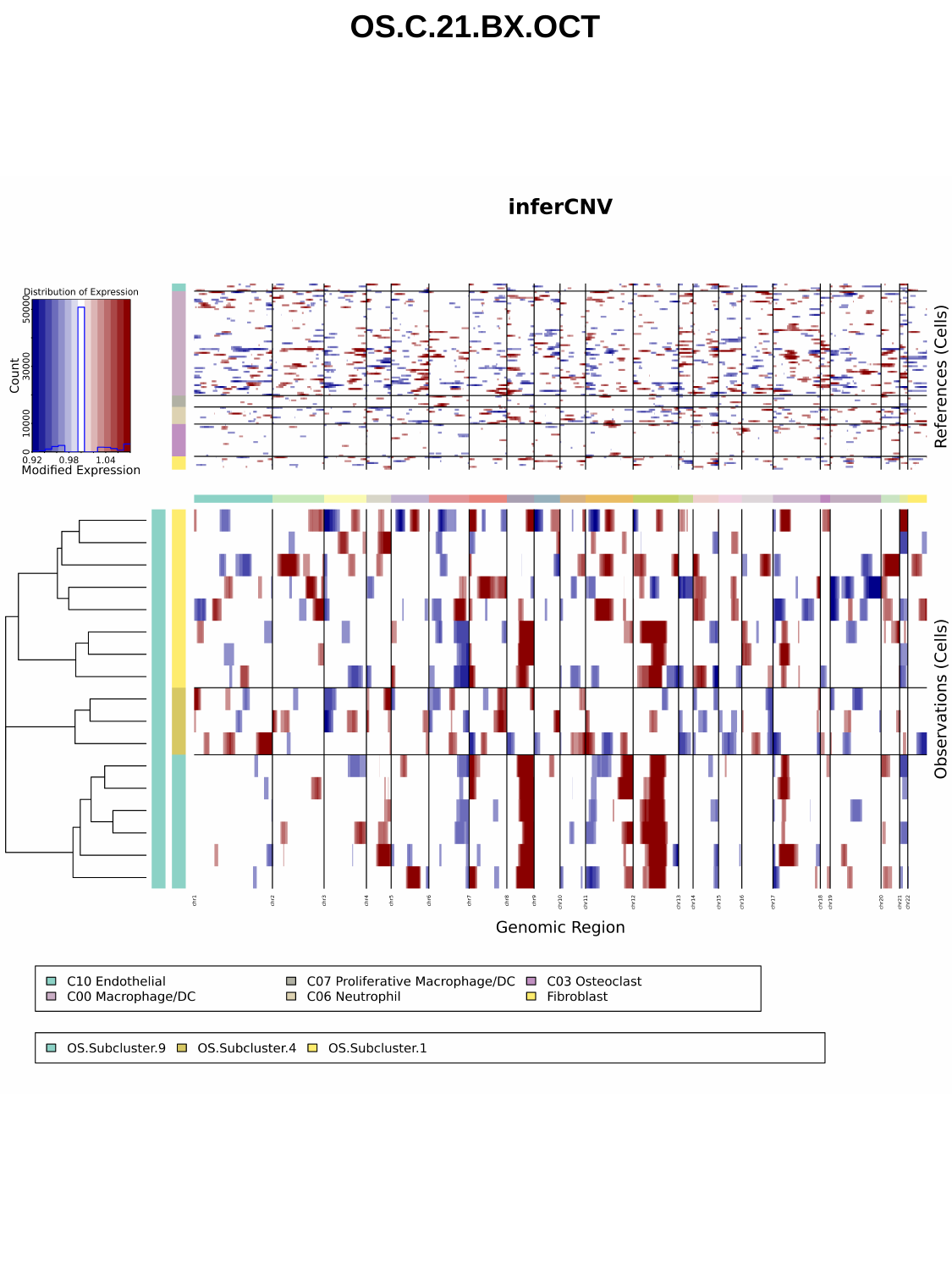

OS.C.21.BX.OCT

### Slide 5
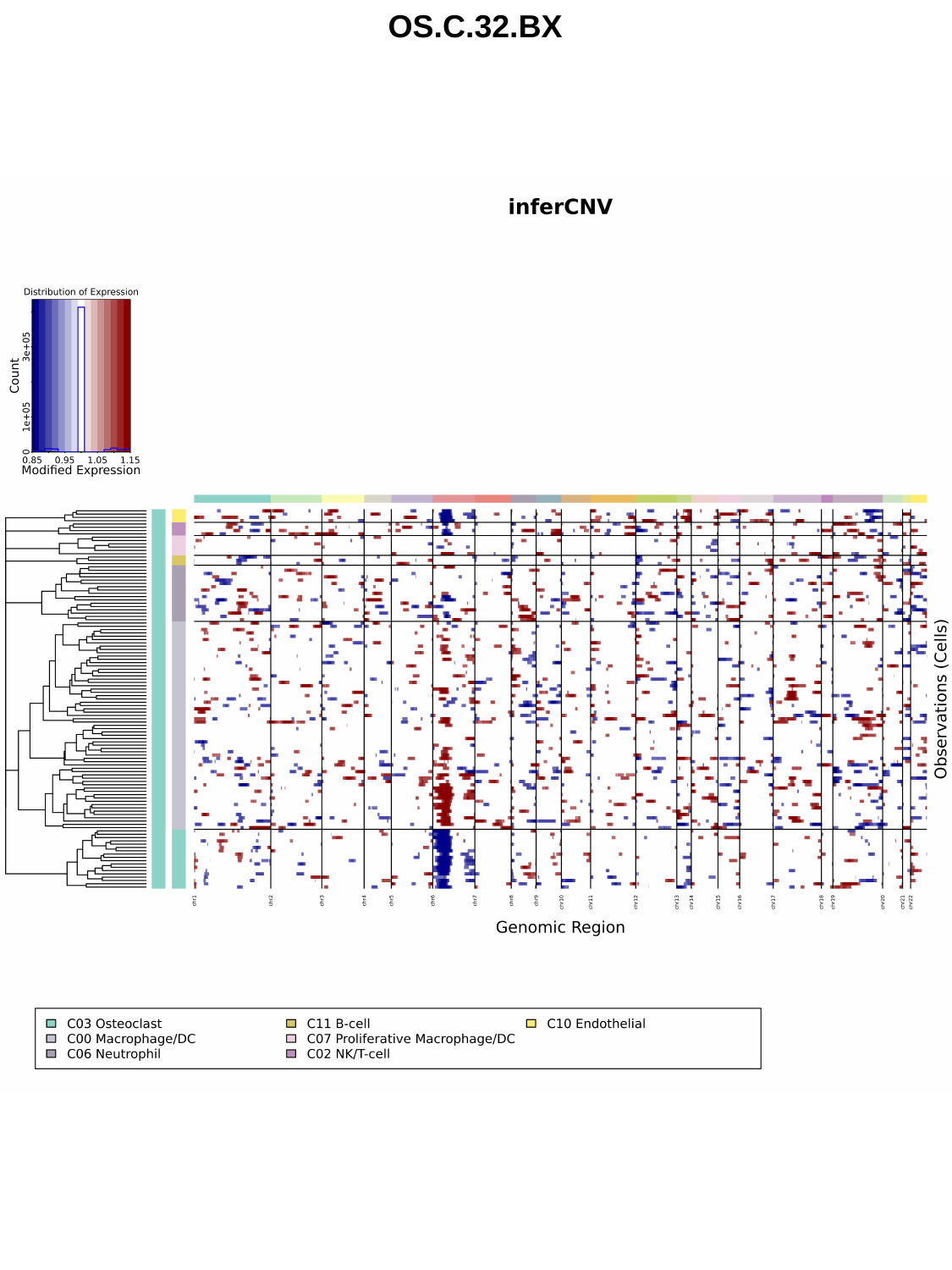

OS.C.32.BX

### Slide 6
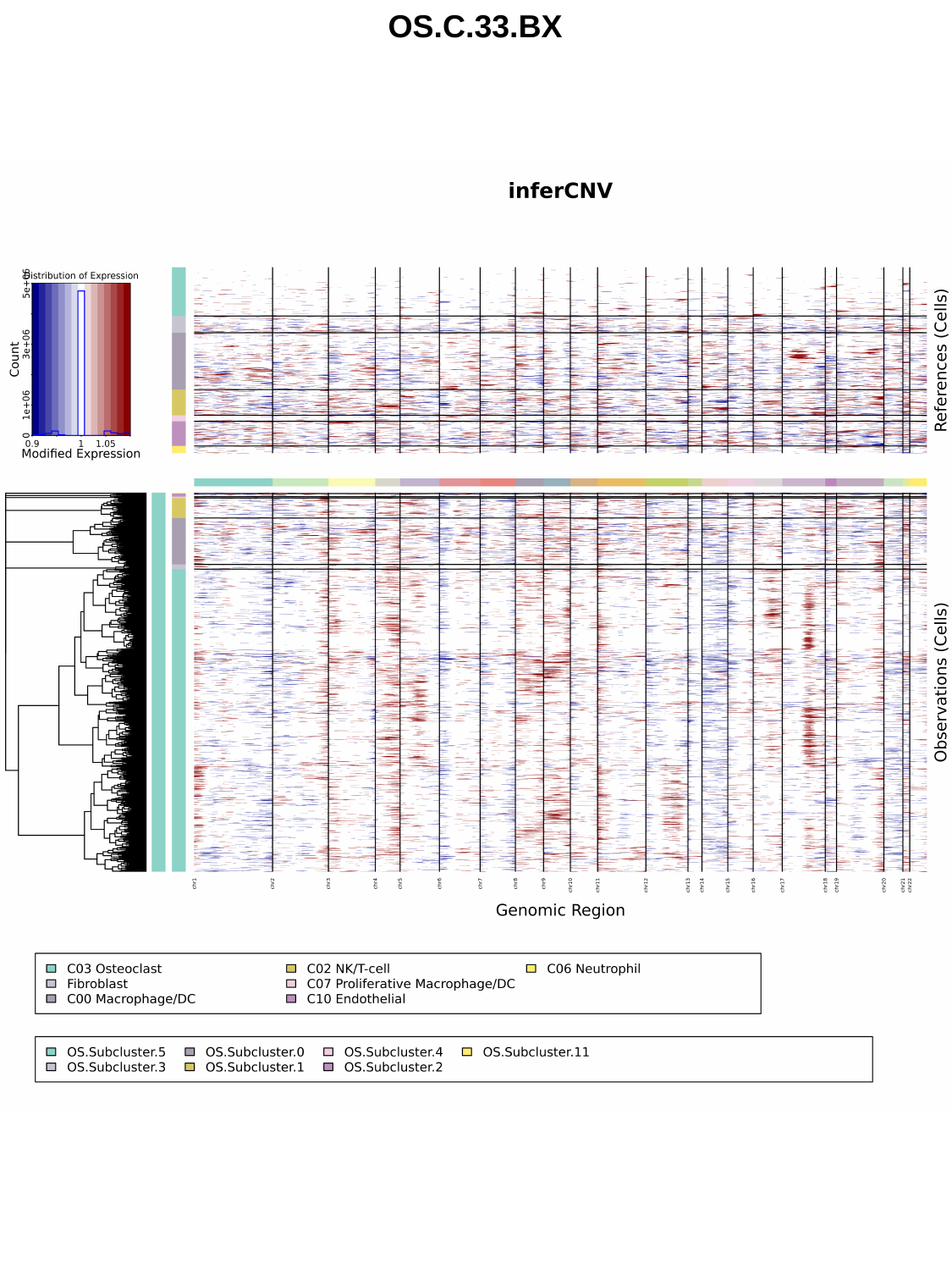

OS.C.33.BX

### Slide 7
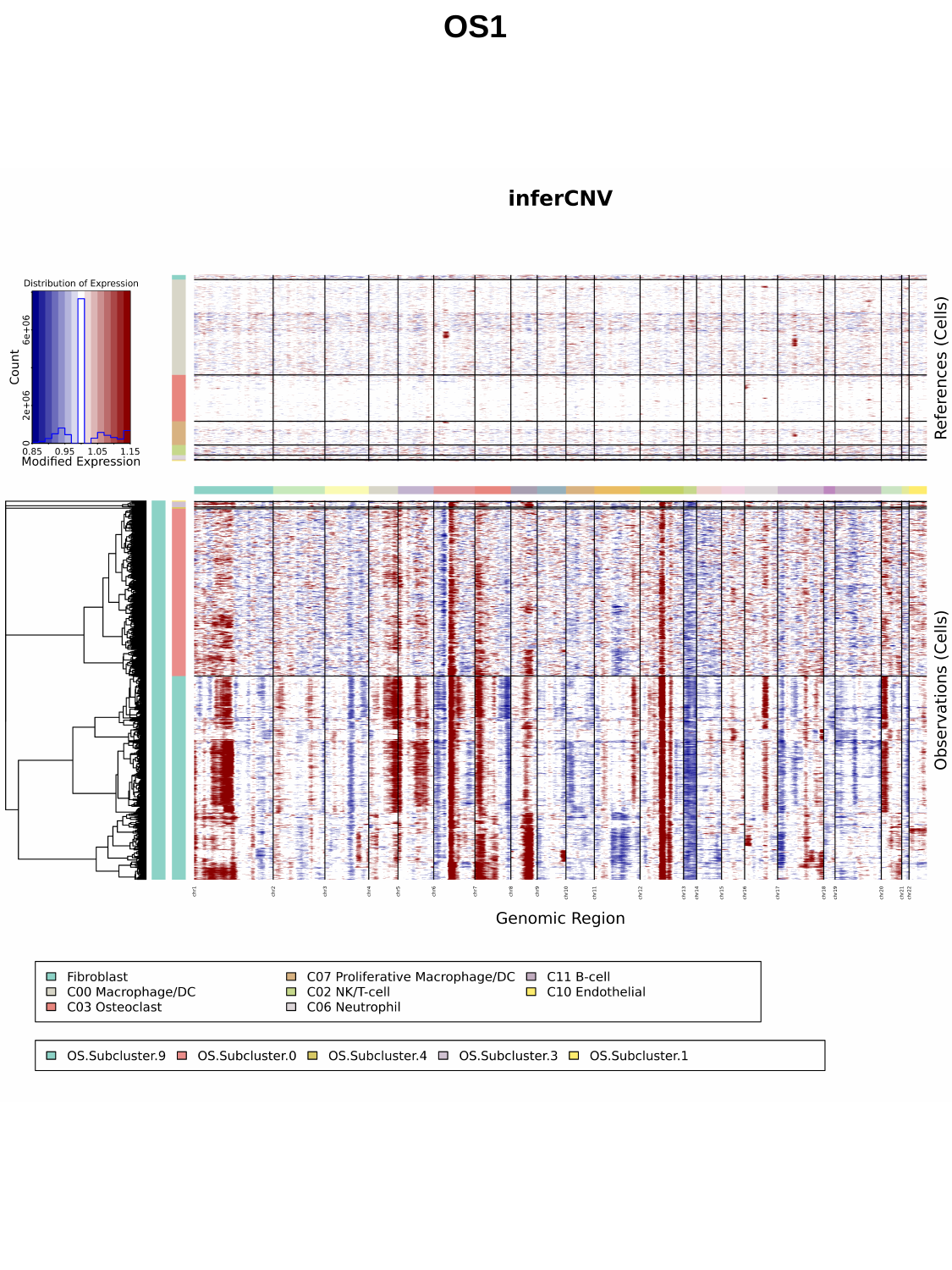

OS1

### Slide 8
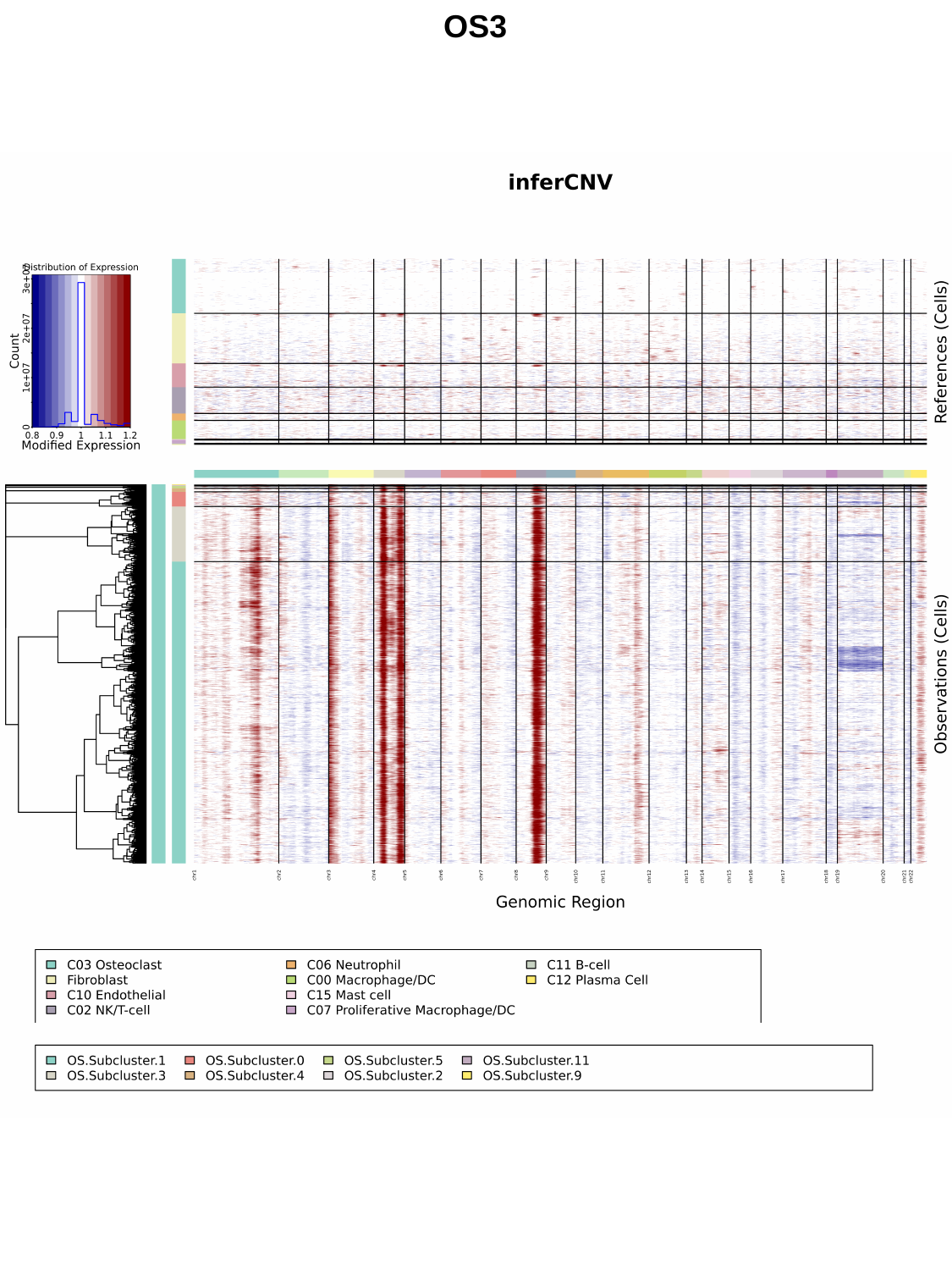

OS3

### Slide 9
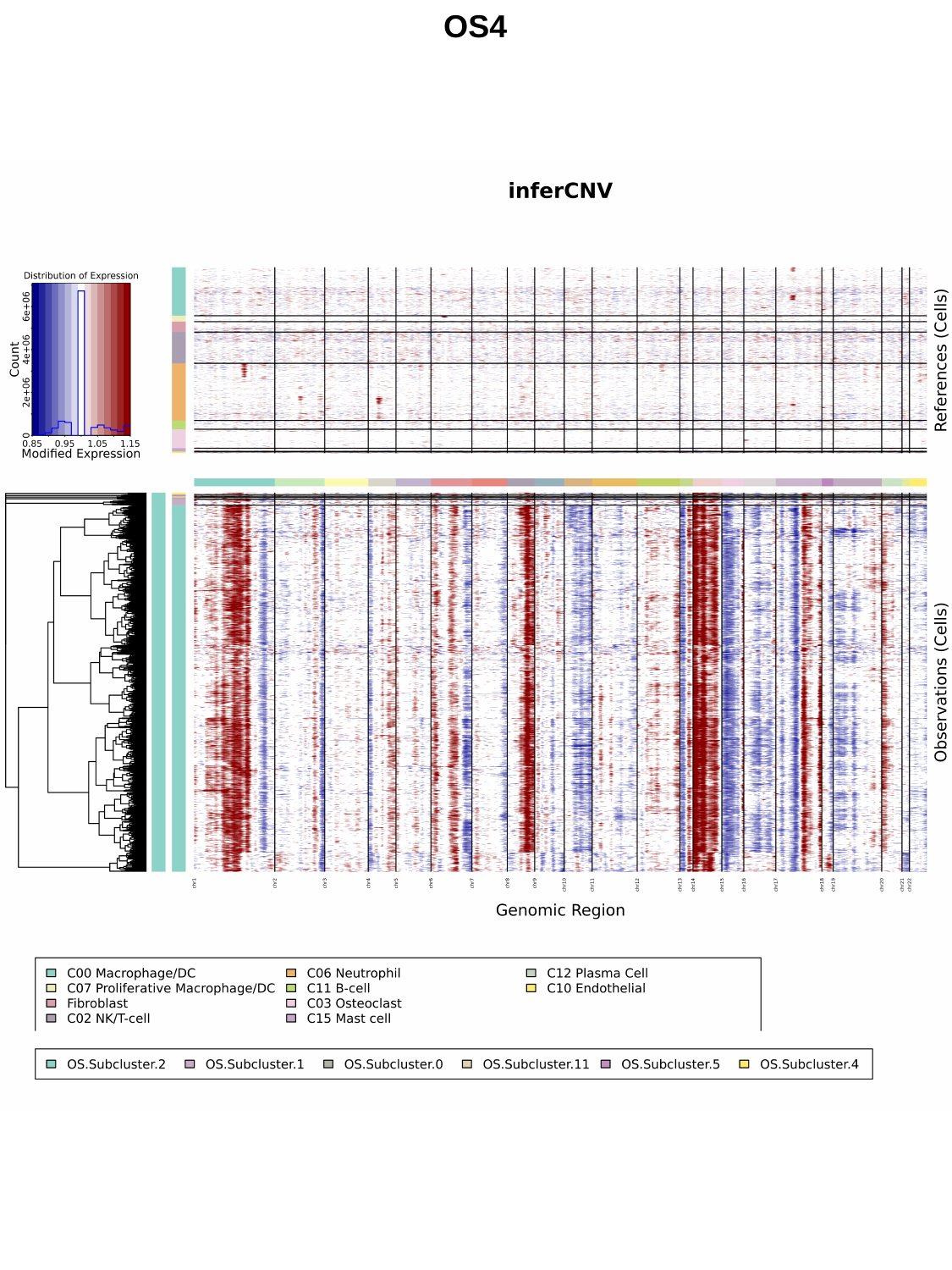

OS4

### Slide 10
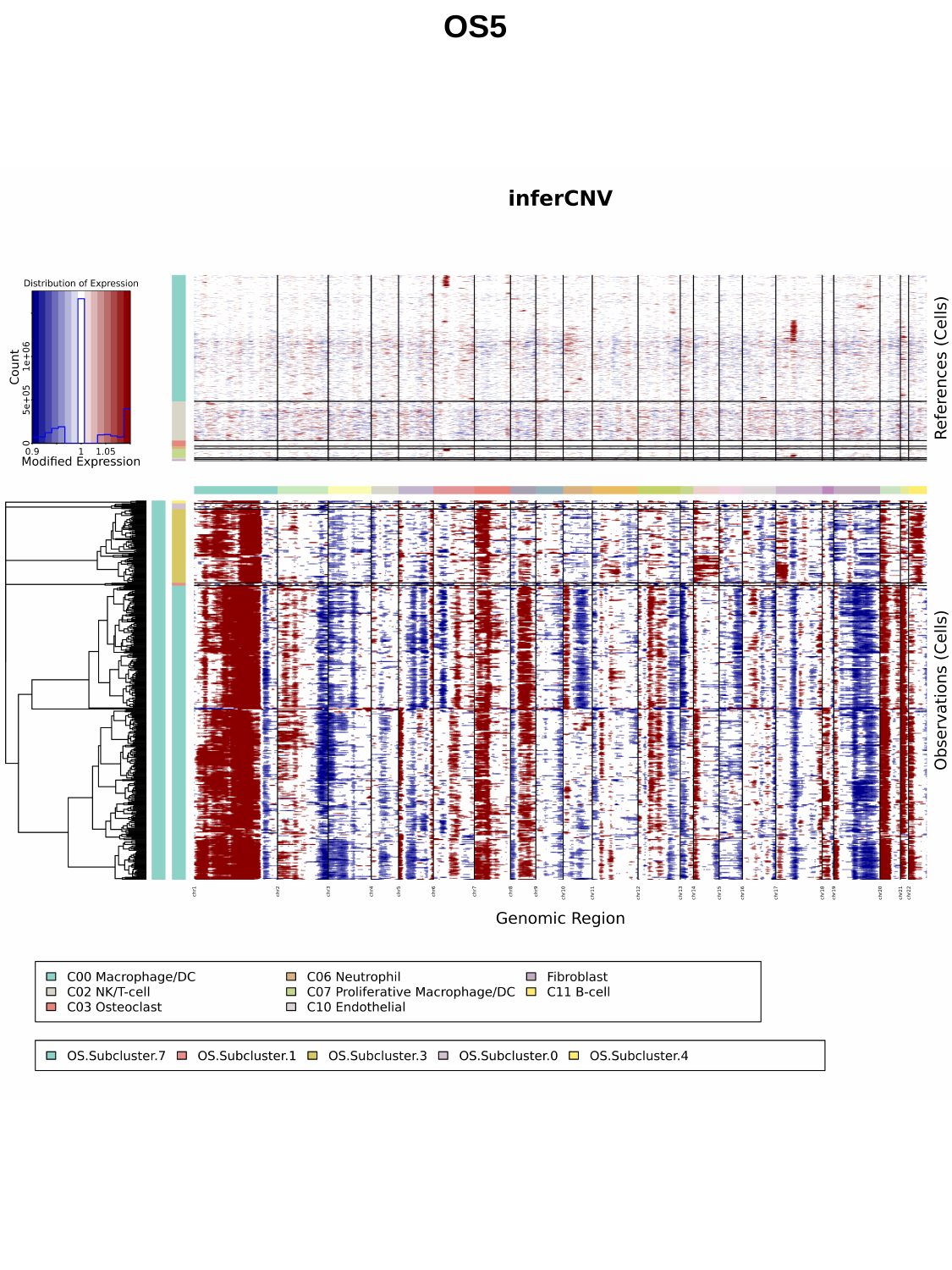

OS5

### Slide 11
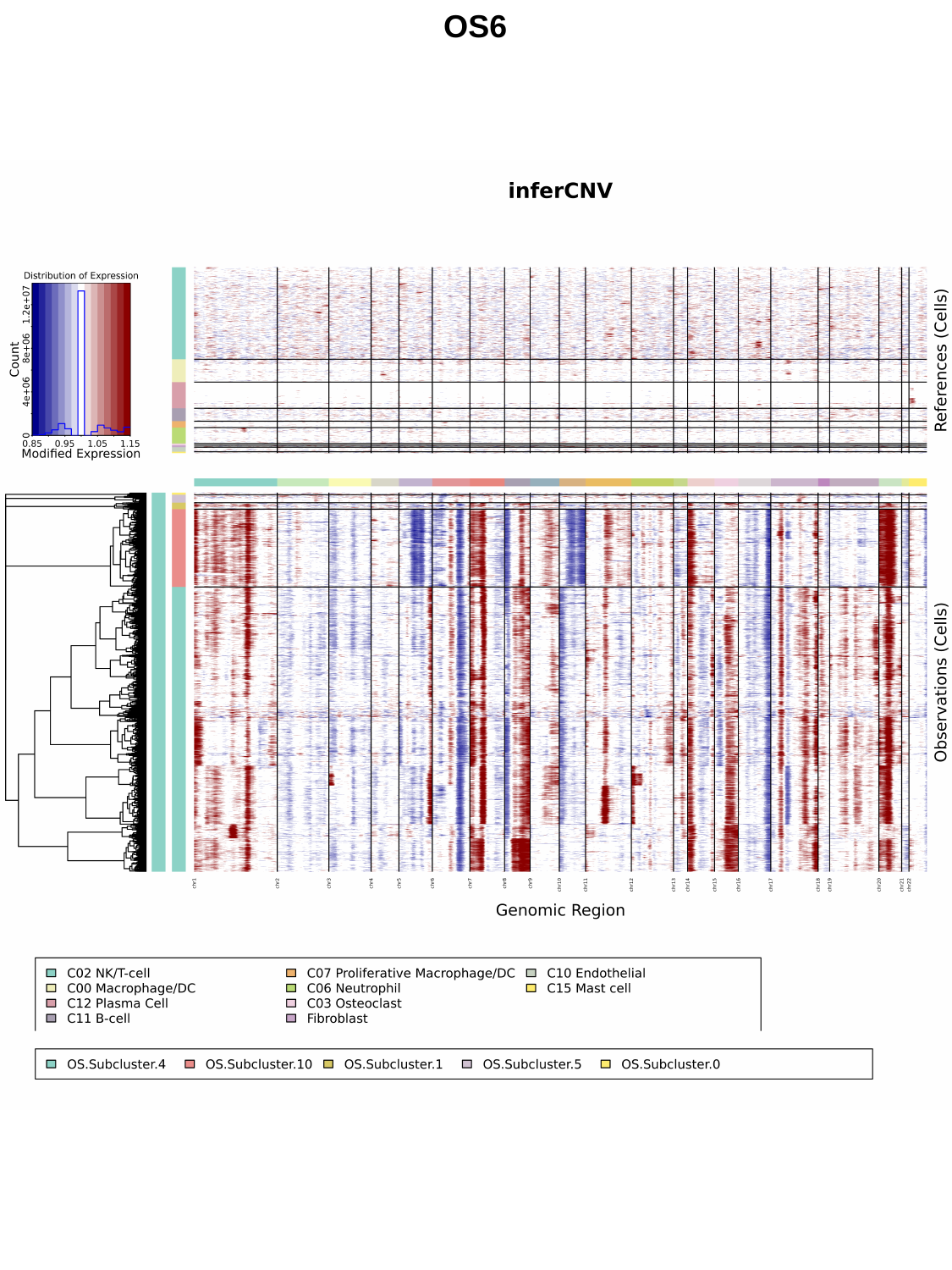

OS6
